# Unusual evolution of tree frog populations in the Chernobyl exclusion zone

**DOI:** 10.1101/2020.12.04.412114

**Authors:** Clément Car, André Gilles, Olivier Armant, Pablo Burraco, Karine Beaugelin-Seiller, Sergey Gashchak, Virginie Camilleri, Isabelle Cavalie, Patrick Laloi, Christelle Adam-Guillermin, Germán Orizaola, Jean-Marc Bonzom

**Affiliations:** Institut de Radioprotection et de Sûreté Nucléaire (IRSN), PSE-ENV/SRTE/LECO, Cadarache, 13115, Saint Paul Lez Durance, France; UMR RECOVER, Aix-Marseille Université, INRAE, centre Saint-Charles, 3 place Victor Hugo, 13331 Marseille, France; Animal Ecology, Department of Ecology and Genetics, Evolutionary Biology Centre, Uppsala University, Norbyvägen 18D, SE-75236 Uppsala, Sweden; Institute of Biodiversity, Animal Health and Comparative Medicine, College of Medical, Veterinary and Life Sciences, University of Glasgow, Glasgow G12 8QQ, United Kingdom; Chornobyl Center for Nuclear Safety, Radioactive Waste and Radioecology, 07100, Slavutych, Ukraine; Institut de Radioprotection et de Sûreté Nucléaire (IRSN), PSE-SANTE/SDOS/LMDN, Cadarache, 13115, Saint Paul Lez Durance, France; IMIB-Biodiversity Research Institute (Univ. Oviedo-CSIC-Princip. Asturias), Universidad de Oviedo, Campus de Mieres, Edificio de Investigación 5^a^ planta, c/ Gonzalo Gutiérrez Quirós s/n, 33600 Mieres-Asturias, Spain; Zoology Unit, Department Biology Organisms and Systems, University of Oviedo, c/ Catedrático Rodrigo Uría s/n, 33071 Oviedo-Asturias, Spain

## Abstract

Despite the ubiquity of pollutants in the environment, their long-term ecological consequences are not always clear and still poorly studied. This is the case concerning the radioactive contamination of the environment following the major nuclear accident at the Chernobyl nuclear power plant. Notwithstanding the implications of evolutionary processes on the population status, few studies concern the evolution of organisms chronically exposed to ionizing radiation in the Chernobyl exclusion zone. Here, we examined genetic markers for 19 populations of Eastern tree frog (*Hyla orientalis*) sampled in the Chernobyl region about thirty years after the nuclear power plant accident to investigate microevolutionary processes ongoing in local populations. Genetic diversity estimated from nuclear and mitochondrial markers showed an absence of genetic erosion and higher mitochondrial diversity in tree frogs from the Chernobyl exclusion zone compared to other European populations. Moreover, the study of haplotype network permitted us to decipher the presence of an independent recent evolutionary history of Chernobyl exclusion zone’s Eastern tree frogs caused by an elevated mutation rate compared to other European populations. By fitting to our data a model of haplotype network evolution, we suspected that Eastern tree frog populations in the Chernobyl exclusion zone have a high mitochondrial mutation rate and small effective population sizes. These data suggest that Eastern tree frogs populations might offset the impact of deleterious mutations because of their large clutch size, but also question the long term impact of ionizing radiation on the status of other species living in the Chernobyl exclusion zone.

## Introduction

The loss of biodiversity during the past 50 years is unprecedented in human history. Pollution, as part of the major drivers of biodiversity loss (namely habitat and climate change, pollution, overexploitation of natural resources, and invasive species), has severely altered many ecosystems^1^. Among the large diversity of pollutants, radioactive contamination caused by human activities, and the associated risks for ecosystems and humans, are the subject of broad societal and scientific concern^2^. This is particularly true in the case of major nuclear accident such as the one occurred at the Chernobyl nuclear power plant (NPP) on April 1986^3,4^. Although the short-term adverse effects of high ionizing radiation doses on wildlife following this accident are not questioned^5-7^, there are still many unknowns and controversies on the long-term ecological consequences of these radioactive releases^8-11^.

One of the biggest challenges for an accurate estimation of the impact of chronic pollution on ecosystems, is to understand, quantify and predict its effects not only at individual, but also at population level^12-14^. Understanding the impact of pollutants on populations allows to investigate evolutionary processes that may affect population status and their capacity to persist in the future. Several studies in the Chernobyl area have estimated the abundance and interspecific diversity of wildlife after the accident ^15-29^ However, these studies have provided inconclusive, and often divergent results, dependent on the sampling design (e.g. for mammals^17,23,30^). In addition, studies investigating the evolution of wildlife in Chernobyl area are scarce, and have not provide solid conclusions ^31,32^ In order to increase our understanding on the impact of ionizing radiation on wildlife in the Chernobyl area, we must examine intraspecific genetic variations. Examining genetic variations within and between populations may allow to estimate differences in the intensity of possible evolutionary processes occurring in wildlife populations^33-36^. Evolutionary processes (mutation, migration, genetic drift, selection) must be understood as the mechanisms at the origin of the modification of genetic variations within populations. Genetic diversity indices, in particular, can be highly informative from an ecological perspective since changes in genetic diversity can affect the capacity of populations to cope with environmental change^37-41^.

Populations exposed to pollutants often experience genetic erosion^35^. Two processes can be at the origin of this decreased diversity: a directional selective pressure which can be driven by the modification of the environment^36,42^, and/or a demographic bottleneck involving the fixation of polymorphic alleles with neutral drift^38,43-45^. Most of the population genetic studies carried out in the Chernobyl area have been conducted on the bank vole *Myodes glareolus,* and showed increased genetic diversity in highly radio-contaminated areas^46-51^. There are two not mutually exclusive explanations for this observation. First, exposure to radioactive pollution can lead to an increased mutation rate^47,52,53^, which can partially offset the genetic diversity loss caused by population bottlenecks. Alternatively, the Chernobyl exclusion zone (CEZ) - which is an area established soon after the Chernobyl nuclear disaster where human population was evacuated^54^ - could act as an ecological sink^55-58^: a demographic deficit caused by the polluted habitat (mortality > natality) could lead to immigration to these habitats, and *in fine* to an increase in genetic diversity^49,59,60^.

Here, we examine the relationship between radionuclide contamination in the CEZ and the genetic pattern of populations of a lissamphibian species, the Eastern tree frog (*Hyla orientalis*)^61^ Bedriaga 1890 (Anura, Hylidae). The phylogeography of this species is well understood which allows the examination of Chernobyl populations in the context of the general evolutionary history of the species^62^. In addition, the Eastern tree frog may be significantly exposed to ionizing radiation in both aquatic and terrestrial environments at susceptible stages of its development, especially during the metamorphosis and during its hibernation in the contaminated soil^63,64^.

We studied population genetics from 19 populations of *H. orientalis* sampled about thirty years after the Chernobyl NPP accident at sites located across a wide range of radioactive contamination inside and outside the CEZ (Fig. 1.b). We used mitochondrial and nuclear genetic markers. These markers differ in their mode of transmission, rate of evolution, and dynamics against environmental disturbances^65-67^. Genetic diversity of populations from the CEZ was compared to that of populations distant up to 40 km from the CEZ (Slavutych), as well as to five other European populations belonging to the same clade^62^ (Fig. 1a). Finally, we studied the mitochondrial haplotype network and made simulation of networks over 10 and 15 generations in order to estimate the population parameters of frogs living in the CEZ since the accident.

**Figure 1:**
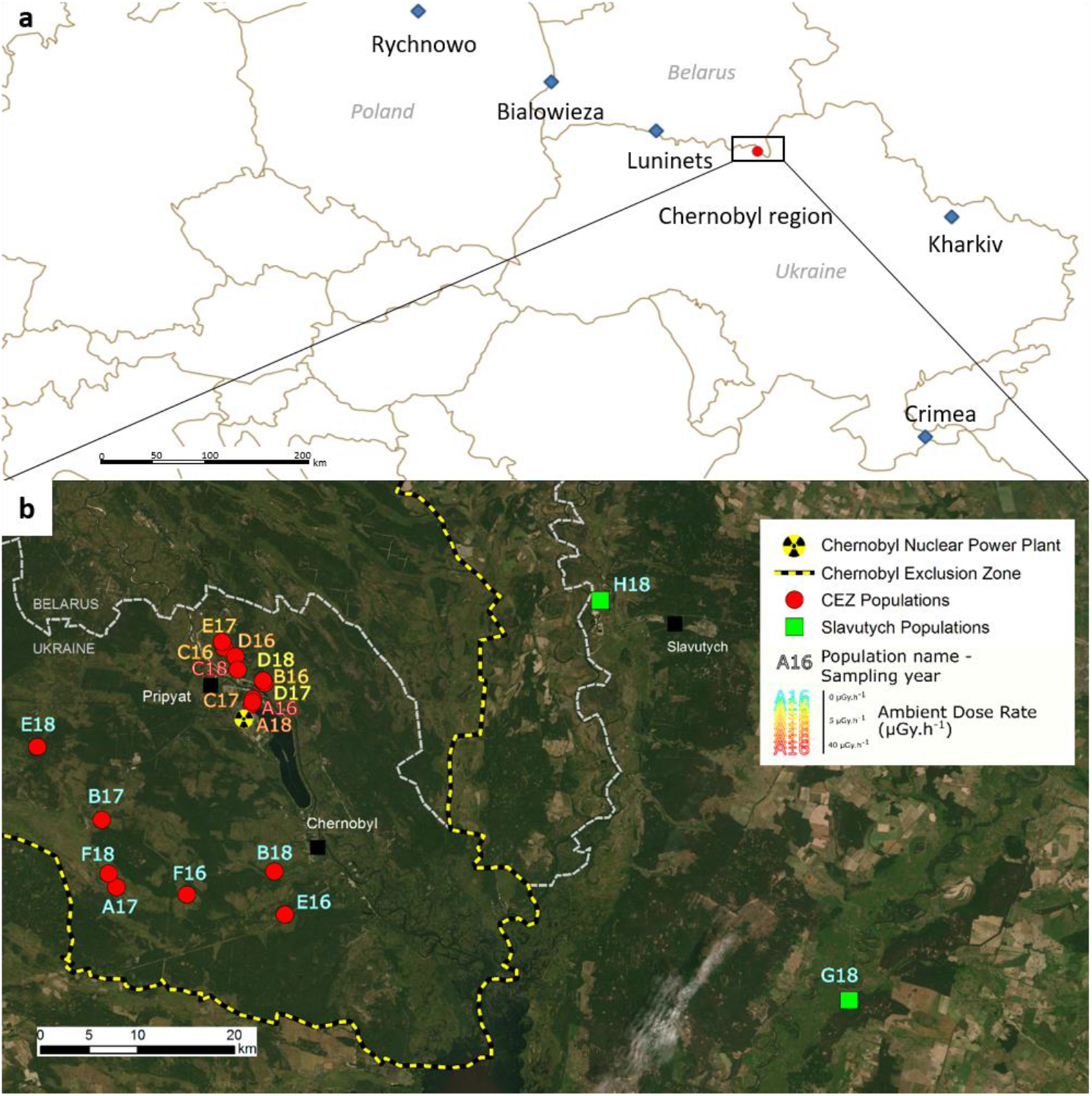
**a**. Location of European populations of Eastern tree frogs outside the Chernobyl region sampled by Dufresnes et al.^62^ (blue diamonds) and the 19 populations sampled at the Chernobyl region (red circles). **b.** Map of the Chernobyl region and location of the 19 populations sampled in 2016, 2017, 2018 in the CEZ and at Slavutych. The map was created with ArcGis v. 10.5. Source and service layer credits for satellite imagery: Esri, DigitalGlobe, GeoEye, Earthstar Geographics, CNES/Airbus DS, USDA, USGS, AeroGRID, IGN, and the GIS User Community.

## Results

### Mitochondrial DNA heteroplasmies

Based on the analysis of the mitochondrial DNA (mtDNA) of 216 Eastern tree frogs *Hyla orientalis* sampled in the CEZ and at Slavutych, we observed 20 substitutions composed of 19 transitions (12 C/T and 7 A/G) and 1 transversion (A/T) (Supplementary Tables 1 and 2). By comparing the haplotypes found in the Chernobyl region with the haplogroups described for other areas of Europe^61,62^, we determined that they were part of the clade D4, characteristic of areas from the northern Black Sea shores to the Baltic Sea. In addition, we detected seven individuals from three populations with two different haplotypes in the CEZ which we considered as being cases of heteroplasmy, while none were detected in the other European populations (see Methods, Supplementary Fig. 1 and Supplementary Table 3).

### High mitochondrial genetic diversity for CEZ populations

We compared the genetic diversity of populations with sufficient sample size (n ≥ 7 individuals) sampled in the CEZ to the populations from Slavutych and to those from other European areas sampled by Dufresnes et al.^62^. Mitochondrial haplotypic and nucleotide diversities of all populations from CEZ (h = 0.7308, π = 0.0024) were significantly higher (W = 91, p < 0.005) than those of other European populations (h = 0.6071, π = 0.0008) (Fig. 2a and Supplementary Table 4). The lowest mitochondrial haplotype diversity was measured for the populations nearest to Chernobyl region, in Kharkiv, Ukraine (h = 0.2500) and Luninets, Belarus (h = 0) (Figure 1a and Figure 2b). The two Slavutych populations present an intermediate mitochondrial haplotype diversity, as the H18 population had low genetic diversity (h = 0.2857), while the G18 population had a genetic diversity fairly close to the genetic diversity of the CEZ populations (h = 0.6444) (Fig. 2b). To compare the variations of mtDNA to nuclear DNA (nDNA) we examined 21 nuclear microsatellites from 126 individuals captured in 2016 and 2017 (Supplementary Table 8). Unlike mtDNA, all the indices of estimated nuclear genetic diversity, such as the genetic diversity within populations (HS) (Fig. 2c), showed no significant differences between the CEZ populations and the other European populations (Supplementary Table 5).

**Figure 2:**
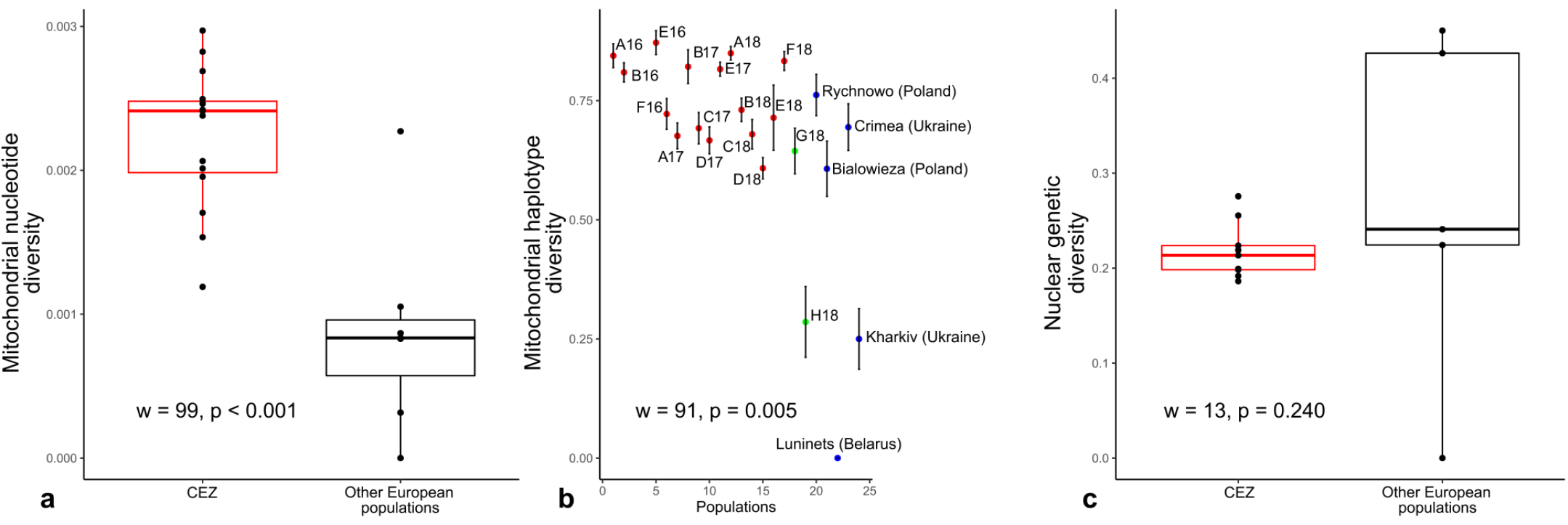
Comparison between genetic diversity estimates at the European level. **a.** Boxplot of mitochondrial nucleotide diversity (i.e. the probability that two randomly chosen nucleotides of the cytochrome b at a homolog position are different^117,118^) for CEZ (red) and other European populations (black). Genetic diversity is higher at the CEZ than at other European populations (Mann-Whitney, w = 99, p = 0.0004). **b.** Mitochondrial haplotype diversity estimates (i.e. the probability that two randomly chosen haplotypes of the cytochrome b are different^117^) ± standard error for CEZ (red), populations from Slavutych (green) and sampled by Dufresnes et al. (blue)^62^. Genetic diversity is higher at the CEZ than at other European populations (Mann-Whitney, w = 91, p = 0.005). **c.** Boxplot of nuclear genetic diversity estimated on the 21 microsatellites markers^117^ for CEZ (red) and other European populations (blue). There are no significant differences between the genetic diversity of CEZ and other European populations (Mann-Whitney, w = 13, p = 0.240).

We also investigated the relationship between genetic diversity and the average total dose rates (ATDRs) of ionizing radiation absorbed by individuals in each population of the Chernobyl region (both inside CEZ and around Slavutych; Fig. 1b, see Methods). ATDRs were estimated to provide a more accurate description of the exposure of populations to ionizing radiation than the generally used ambient dose rate, by taking into account the contribution of the different radionuclides and radiation types (alpha, beta and gamma emitters) from all exposure pathways (internal and external) (See Giraudeau et al.^63,68^ as well as Methods and Supplementary note 1). Only mitochondrial nucleotide diversity was significantly positively correlated to ATDRs (S = 294, rho = 0.640, p = 0.007) (Fig. 3a and Supplementary Table 6). In contrast, the correlation between mitochondrial haplotype diversity and ATDR was not significant (S = 658, rho = 0.193, p = 0.455; Fig. 3b). Genetic diversity in nuclear microsatellites was not significantly correlated with ATDRs, although these parameters showed a non-significant negative correlation (S = 194, rho = −0.617, p = 0.086; Fig. 3c). Only private allelic richness and ATDRs were significantly negatively correlated (S = 221.13, rho = −0.843, p = 0.004) (Supplementary Table 7).

**Figure 3:**
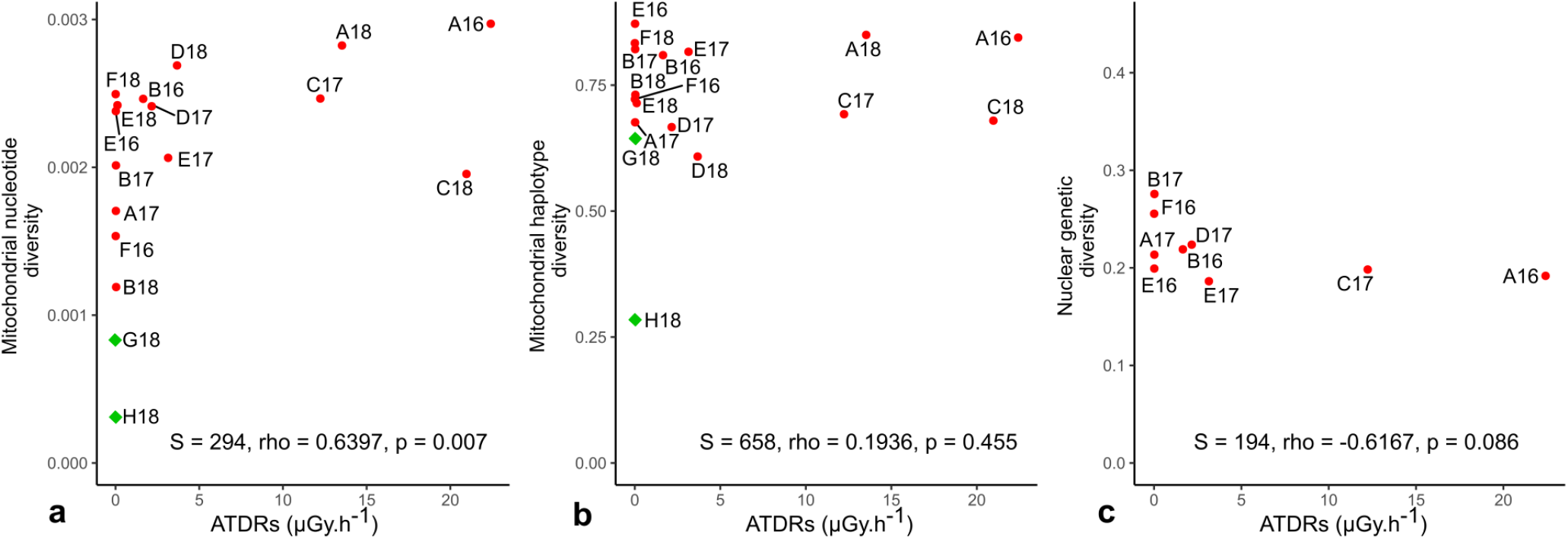
Correlation plots representing genetic diversity estimates on population-averaged dose rate (ATDR) in μGy.h^-1^. Only populations of the Chernobyl region (i.e. CEZ (red dots) and Slavutych (green diamonds), Fig. 1b) with sample size > 7 individuals were compared. **a.** Mitochondrial nucleotide diversity estimates (i.e. the probability that two randomly chosen nucleotides of the cytochrome b at a homolog position are different^117,118^) on ambient dose rate of the corresponding population. Nucleotide diversity is positively correlated to ATDR (S = 294, rho = 0.6397, p = 0.007). **b.** Mitochondrial haplotype diversity estimates (i.e. the probability that two randomly chosen haplotypes of the cytochrome b are different^117^) on ATDR of the corresponding population. Haplotype diversity is not correlated to ATDR (S =, 658 rho = 0.1936, p = 0.455). **c.** Nuclear genetic diversity (Hs) estimated on the 21 microsatellites markers^117^ on ATDR. Genetic diversity is not correlated to ambient dose rate (S = 194, rho = −0.6167, p = 0.086).

### Local geographical structure of genetic variation

In order to study the genetic structure of populations sampled in the CEZ and the Slavutych area, we used a pairwise genetic differentiation estimator between populations (pairwise Fst)^69^. Unlike the low differentiation estimated with nuclear microsatellites (−0.031 < Fst < 0.093), differentiation estimated from cytochrome b sequences was relatively high (−0.173 < Fst < 0.426). The highest genetic differentiations for mitochondrial marker (cytochrome b) were observed between the South West A17 and North D18 populations, and other populations (0.072 < Fst < 0.426, Fig. 4a). Slavutych populations were also highly genetically differentiated from CEZ populations (0.095 < Fst < 0.395). Genetic pairwise differentiations estimated on nuclear markers were similar to those estimated on mitochondrial markers, the most differentiated population being A17 (0.021 < Fst < 0.093, Fig. 4b). Despite the absence of a complete similarity between geographical and genetic structures (Figs. 1b and 4a,b), the genetically closest populations were, as expected, usually the geographically closest populations. This similarity was obvious when separating populations in Neighbour Joining (NJ) trees build for each sampling year (Supplementary Fig. 2).

**Figure 4:**
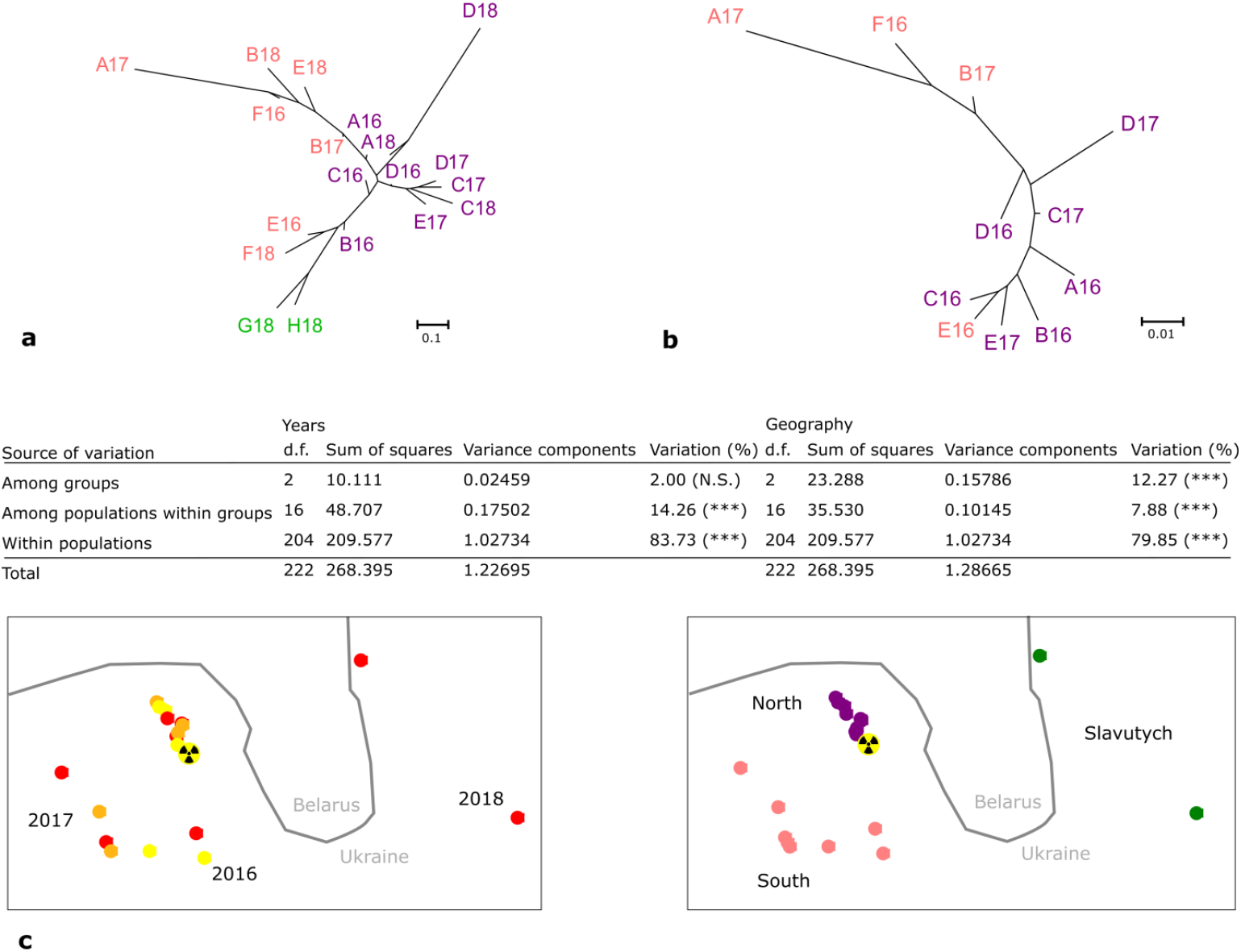
Genetic structure of the 19 populations of Eastern tree frogs from CEZ and Slavutych. Neighbor-joining tree were constructed from genetic distances calculated as Fst_posi_/(1-Fst_posi_) with Fst_posi_ equal to the addition of Fst and the absolute value of the lowest Fst in order to avoid negative values and respect proportionality of pairwise Fst (see Methods for details). **a**. Neighbor Joining tree of CEZ (purple and pink) and Slavutych (green) populations from cytochrome b (mtDNA). **b**. Neighbor-Joining tree of CEZ populations (red) from microsatellites (nDNA). **c.** AMOVA analysis conducted on Year and Geographical groups on mtDNA. Stars represent significance calculated from Arlequin with 1023 permutations^121^ (***: sign < 0.001). Year groups are 2016, 2017, 2018 (2016: yellow, 2017: orange, 2018: red) and geographical groups are north close to the Chernobyl Nuclear Power Plant (radiation warning symbol), south distant form the north and Slavutych (north: purple, south: pink, Slavutych: green).

We examine the effects of year of sampling, and sampling site on the distribution of mitochondrial genetic structure, using Analysis of Molecular Variance (AMOVA). We used three year groups (2016, 2017, 2018), and three geographical areas corresponding to three groups of populations in the Chernobyl region: one in the North close to the NPP, one on the South of the exclusion zone, and one including the Slavutych populations (Fig. 4c). In both cases, the highest variance was observed within populations (83.75% and 79.85%). However, the inter-group variance based on years was not significant (2.00%, p > 0.05), in opposition to the variance based on geographical regions (12.27%, p < 0.001).

In order to test if the increase of genetic distance between populations was shaped by their geographic distances (“isolation by distance” hypothesis), we run Mantel test^70^ between pairwise genetic distances matrix estimated from Fst and pairwise geographic distances matrix. Isolation by distance was significant for both nuclear (r = 0.4453, p = 0.005), and mitochondrial markers (r = 0.3461, p = 0.009). The correspondence between nuclear and mitochondrial genetic distances described from NJ trees was also significant (r = 0.6627, p = 0.003). Because of a possible link between geography and radionuclide contamination^71^, we also tested if isolation by distance was carried by the ATDRs using a partial Mantel test. The correlation between genetic distances and geographic distances could not be explained by the differences of ATDRs (mtDNA: r = 0.3474, sign = 0.007; nDNA: r = 0.4452, sign = 0.006). These results indicate that the local genetic structure was mainly influenced by geography, but not by years of sampling or levels of radiation exposure.

### Haplotype networks and CEZ independent evolutionary processes

Following the first quantitative part of our study, we then focused on cytochrome b mitochondrial marker as it allowed us studying qualitatively haplotypes of all populations (i.e. CEZ, Slavutych, and populations outside Chernobyl region sampled by Dufresnes et al.^62^; see Fig. 1a), and examined their genealogical links within a network of haplotypes. This approach is a good way to situate populations within an evolutionary context and explore more subtle evolutionary processes than with diversity indices only^56^. By comparing the haplotype network of the Chernobyl populations (CEZ and Slavutych areas) with the haplotypes of populations outside the Chernobyl region, we identified a single haplotype common to all populations, the central haplotype (Fig. 5). Because of the star-like distribution of the haplotype network of populations outside the Chernobyl region (in blue, Fig. 5) with respect to this central haplotype, we considered it as the ancestral haplotype.

**Figure 5:**
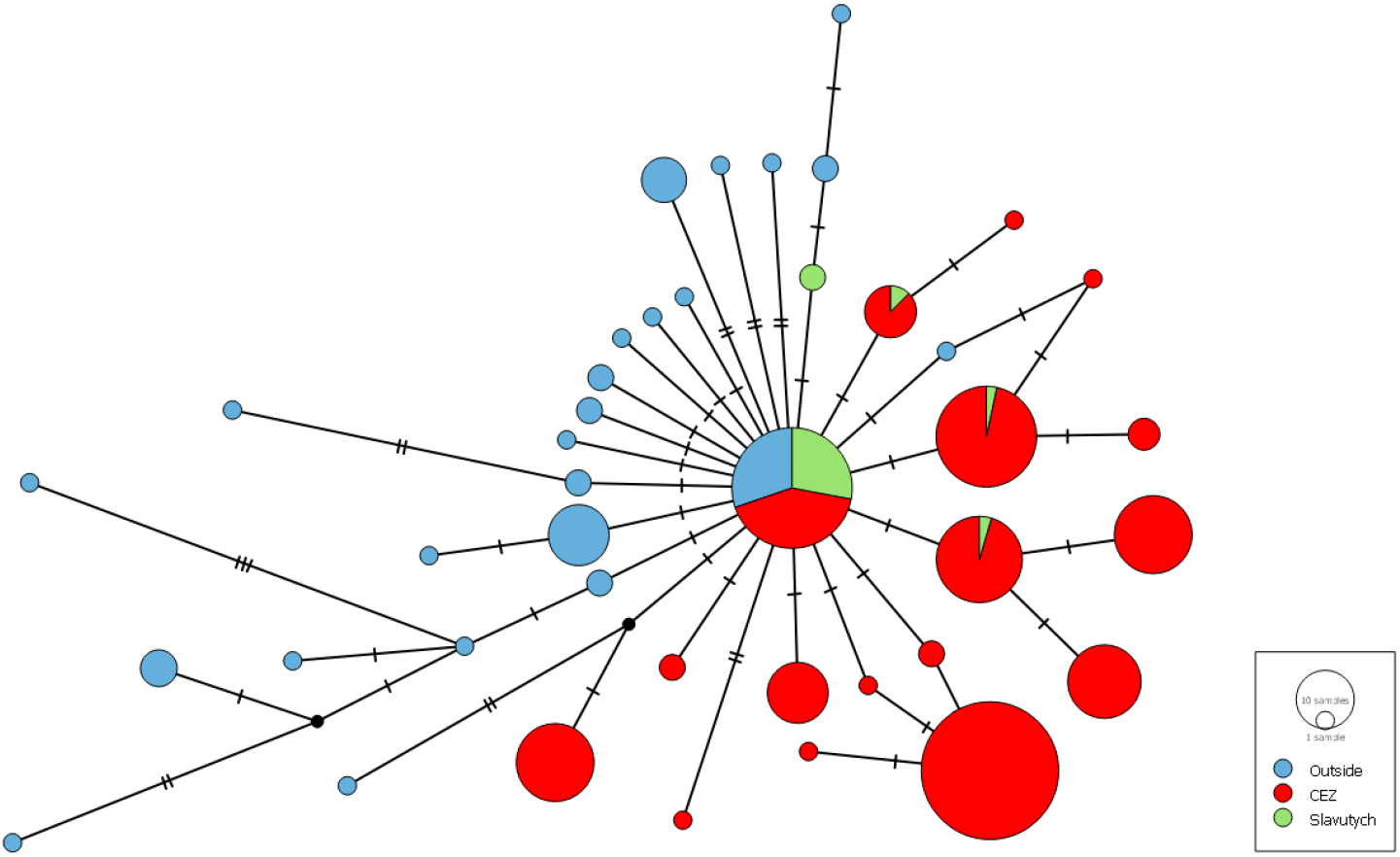
Haplotype network constructed for Eastern tree frog cytochrome b sequences from CEZ (red), Slavutych (green) populations, and European populations sampled by Dufresnes et al.^62^ (blue) using the Median-Joining method^126^ and POPART software^127^. Circles representing haplotypes, their diameter is proportional to the number of individuals and the number of horizontal bars between haplotypes representing the number of nucleotides differing between haplotypes. The network structure can inform on the demographic status of populations: when the central haplotype is large compared to the surrounding haplotypes and lot of one step rare haplotypes surround this central haplotype (e.g. Slavutych and European populations), the population is in demographic expansion; if the central haplotype is not mainly represented and if there are a lot of two or three steps large haplotypes, the population is at the equilibrium mutation/drift and is often formerly diversified (CEZ populations).

We detected a discrepancy between the structure of the CEZ haplotype network and those of all other populations, since the population sampled in Slavutych segregated similarly to populations from other European areas analysed by Dufresnes et al.^62^: the largest haplotype was the central haplotype, surrounded by many one substitution step rare haplotypes (Fig. 5; green and blue). These populations outside the CEZ are in demographic expansion, as confirmed by the rejection of the equilibrium mutation/drift hypothesis (D Tajima = −2.2180, p < 0.01; Fu and Li D* = −4.4028, p = 0.002; R2 = 0.0289, p = 0.001). In contrast, the CEZ populations present a different pattern represented by haplotypes at one and two steps from the central haplotype, shared by many individuals (Fig. 5, in red), and these populations are not in demographic expansion (D Tajima = −0.5641, p = 0.332; Fu and Li D* = −1.4653, p = 0.089; R2 = 0.0663, p = 0.357). These results suggested an independent evolutionary history of the CEZ populations compared to other sampled populations, even in the neighbour population of Slavutych.

### Small populations and elevated mutation rate in the CEZ

To decipher the factors shaping the particular pattern of the CEZ haplotype network, we simulated the evolution of its mitochondrial haplotype network using a range of parameters, starting with mitochondrial haplotype diversity of Slavutych populations, corresponding to the G18 or H18 haplotype frequencies (see Methods and Supplementary note 2). Simulations were made for different prior parameters and a set of statistics was chosen to describe the haplotype network at the 10^th^ and 15^th^ generation, corresponding to a generation time of three and two years, respectively. During a first simulation part, haplotype networks were simulated based on a fluctuating population size since the accident in the range of uniform distribution U(1000-5000) and a classical rate of nucleotide substitution in mitochondrial DNA for amphibians of 20.37×10^-9^ per nucleotide per generation^72^ (Supplementary Table 12). The Principal Component Analysis (PCA) was used to compare simulated and observed statistics, and showed that simulated haplotype networks did not match the observed one, the closest Euclidean distance being 11.13, indicating that the diversity of CEZ populations cannot be obtained with this first set of parameters (Fig. 6a.c and Supplementary Fig. 7).

**Figure 6:**
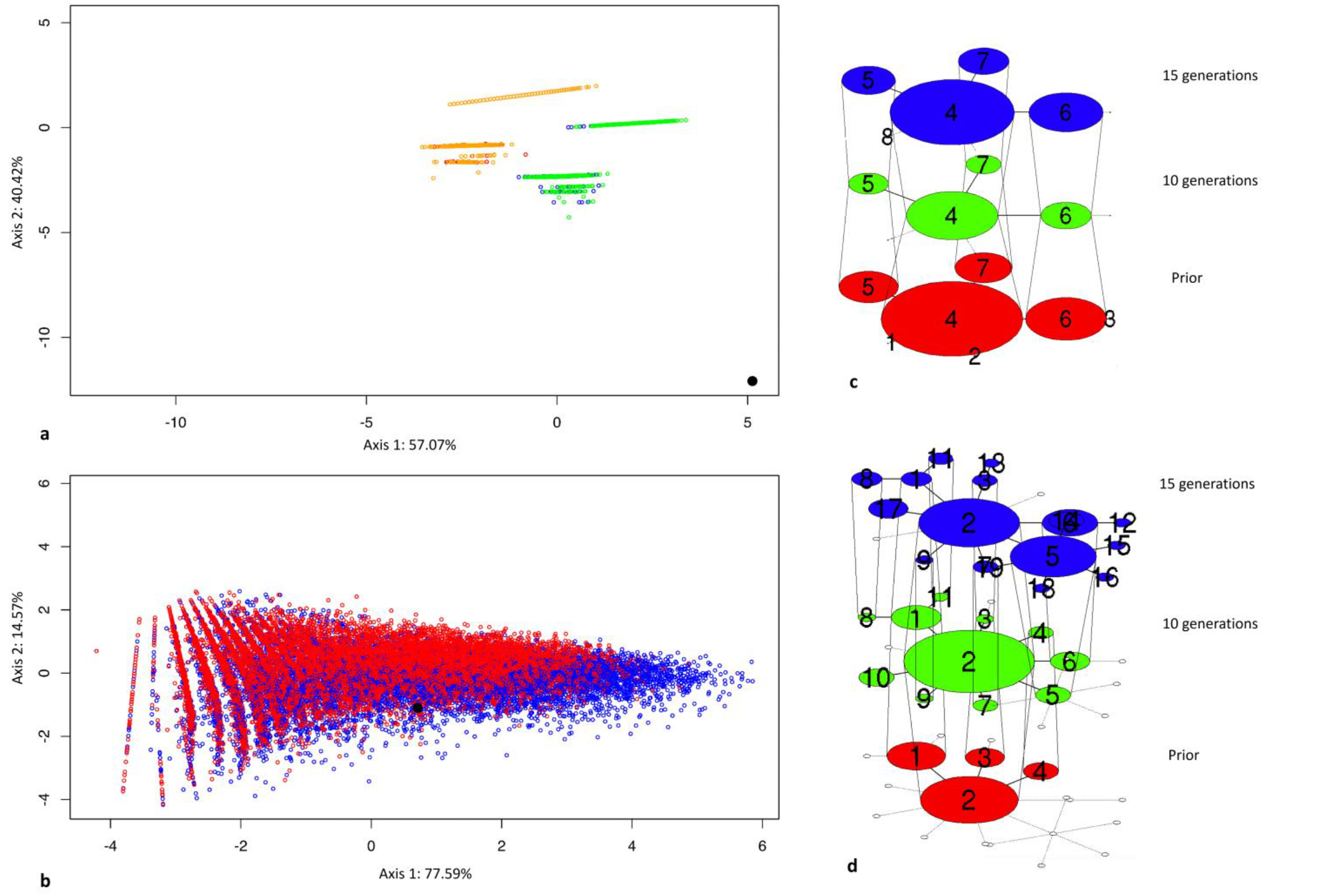
Mitochondrial haplotype network simulation. **a.** Representation of observed (black circle) and simulated (coloured open circles) data with a classical amphibian mitochondrial nucleotide substitution rate (20.37×10^-9^ per nucleotide per generation), a population size sampled in a uniform distribution U(1000-5000), and starting from the H18 (orange) or G18 (green) population haplotype frequencies on the two first axis of a PCA made on a set of haplotype network statistics. The observed data is not in the space of the simulated data. **b.** Representation of observed (black circle) and simulated (coloured circles) data with a high mitochondrial nucleotide substitution rate (0.005, 0.01, 0.02, 0.04, 0.06, 0.08) and small population size (sampled in a uniform distribution U(50,100), U(100,200), or U(200,300)) for 10 (red) and 15 (blue) generations on the two first axis of a PCA made on a set of haplotype network statistics. The observed data is in the space of the simulated data. **c.** One example of haplotype network evolutionary scenario of a simulated population starting from G18 population haplotype frequencies as prior with a classical amphibian substitution rate and populations sizes in the range of uniform distribution U(1000-5000) **d.** One example of haplotype network evolutionary scenario of a simulated population starting from G18 population haplotype frequencies as prior with a high substitution rate (0.04) and a maximal effective size of 100 (for prior (red) and 10 (green) and 15 (blue) generations after the Chernobyl nuclear power plant accident).

In a second simulation of haplotype networks we used smaller population sizes (three modalities of uniform distribution: U(50,100), U(100,200) and U(200,300)) and high nucleotide substitution rates (six modalities: 0.005, 0.01, 0.02, 0.04, 0.06, 0.08 per haplotype per generation in an infinite model site) (Supplementary Table 13). In contrast to the first simulation, PCA displayed a match between the haplotype networks statistics of the simulated and observed data. Indeed, the observed data was in the space of simulated data based on the two first principal components supporting around 90% of variance (Fig. 6b). Posterior parameters were then estimated using a Ward hierarchical cluster analysis on Euclidean distance selecting the 5^th^ percentile of the simulated descriptive statistics closest to the observed descriptive statistics (Supplementary Fig. 8). The closest distance was 0.52 and the median of descriptive statistics for the 5 percentile closest simulated values presented important similarity with observed values (Supplementary Fig. 9). Considering the posterior parameters estimated, the diversity of CEZ populations and the particular haplotype network (Fig. 6d) for the studied mitochondrial marker can thus be obtained in 15 generations with a small population (N_max_ = 100) and a high nucleotide substitution rate of 0.04 per haplotype per generation.

## Discussion

Several studies have shown that in the CEZ, where all residents have been evacuated, large mammals in particular are reappearing doubtless due to a decrease of human disturbance to wildlife ^17,19,20,28^. Conversely, other studies have shown a decrease in the abundance of some species in the CEZ (birds^73^, insects^21^, mammals^23^). There is still no consensus about the long term consequences of the Chernobyl NPP accident, and the effects of exposure to ionizing radiation on population status remain mostly unknown. To date very few studies have focused on the evolutionary processes occurring in natural populations that underwent chronic exposure since the 1986 Chernobyl NPP accident. To the best of our knowledge, our study is the first in the Chernobyl region (i) investigating the evolutionary processes of CEZ populations, in comparison to the global European evolution of the closest lineage to which they belong, (ii) using both qualitative and quantitative mitochondrial genetic information and quantitative nuclear genetic information to estimate the best evolutionary scenario responsible of the observed pattern.

### A higher mtDNA diversity in the CEZ driven by mutation process

In contrast to the expected genetic erosion induced by wildlife exposure to a pollutant^35^, our results did not show a genetic bottleneck of *H. orientalis* populations in the CEZ compared to the other European populations studied by Dufresnes et al.^62^. We found a higher mitochondrial genetic diversity for the populations in the CEZ, while similar nuclear genetic diversity was observed between CEZ populations and other European populations. These results on mitochondrial diversity agree with the increased mitochondrial genetic diversity observed on bank voles, *Myodes glareolus*, from the most contaminated areas of the CEZ^47^. A higher diversity can be explained by two evolutionary processes: migrations from multiple distinct and distant populations, or a local higher mutation rate. The discrepancy between nuclear and mitochondrial markers may orienting towards one of these two mechanisms^74,75^. Indeed repair mechanisms in mtDNA are usually considered less effective than in nDNA^76,77^ notably because of variations in replication mechanism (i.e. low fidelity of the DNA polymerase γ) and a higher number of genome replications per generation especially during oocyte maturation^78^. Thus, the emergence of a mutagenic factor in the environment can induce mutations on mtDNA without increasing nDNA mutations at the same rate. A high migration rate of animals towards the CEZ would increase both mitochondrial and nuclear diversity, a pattern that does not corresponds with our observations. Hence, an increased mutation rate in the CEZ is the most likely explanation to the local genetic novelty and increased genetic diversity for mtDNA and not for nDNA.

### A genetic structure consistent with a higher mutation rate in the CEZ

Mitochondrial and nuclear markers differ also in their range of differentiations between populations, but not in the relative structure of these populations. Indeed, based on pairwise Fst values, the most differentiated populations using mtDNA markers are highly differentiated (> 0.4), but not when using nDNA (< 0.1). The general structure of these populations is quite similar within the CEZ between mitochondrial and nuclear markers (Fig. 4a.b), and for the two type of markers, isolation by distance is not rejected. In amphibians, dispersion is usually male-biased (reviewed by ^79^, but see^80^). Since mtDNA is transmitted by females, in case of a strong migration process, there would have been a discrepancy between the relative nuclear and mitochondrial population genetic structure. These results, thus, confirm the absence of a strong tree frog migration process coming from outside the CEZ, and reaffirm the role of mutation processes occurring on mtDNA. The presence of mitochondrial haplotypes exclusive to the CEZ – in contrast to previous studies on bank voles^81^ – and the absence in the Chernobyl region of haplotypes shared with populations outside the Chernobyl region (except ancestral haplotype), support also the hypothesis of absence of numerous long migration between CEZ and other areas. In this way, the mutation/drift balance explains the higher differentiation found in mtDNA population structure.

### Substitution rate and population size at the origin of a “refugia-like” population

The mitochondrial haplotype network of the CEZ tree frog populations, showed a striking structure that differs from what can be expected from the global demographic expansion of the clade D4^61,62^. This structure is similar to an ancient diversified population, demographically stable even during the last glacial maximum^82,83^. However, it is unlikely that the Chernobyl region would have acted as a refuge zone regarding the global evolutionary history of the *Hyla orientalis* species^62^ and the possible recent impact just after the 1986 Chernobyl nuclear accident on amphibians^5,84^. The results of our simulation suggest that a strong mutation rate coupled with populations of small sizes might be responsible for the establishment of the CEZ haplotype network structure. Our haplotype network simulation obtained the observed CEZ haplotype network pattern in 30 years from control local populations identifying two important parameters, a strong nucleotide substitution per haplotype per generation of 0.04 and populations of small effective size inferior to 100 individuals (Fig. 6). We noticed a better match between the observed network and the simulated network after 15 generations than after 10 generations. Although *H. orientalis* females usually start to breed at 3-year age^85,86^ (i.e. 10 generations from the accident), in the CEZ, female tree frogs may start to breed at 2-year age in order to speed up life-history strategy. A shorter generation time may be an adaptive response to cope with the accumulation of damage in stressful environments^87,88^, as those with radioactive contamination.

### Is ionizing radiation at the origin of an increased substitution rate in the CEZ?

The mitochondrial evolutionary pattern of the CEZ populations, which seems to be the result of a dynamic comparable to an accelerated evolution, is not observed outside the CEZ.

Slavutych’s tree frog populations that are geographically close to the CEZ populations do not show the same haplotype structure and did not present any case of heteroplasmy, contrary to the CEZ populations. Knowing the mutagenic ability of ionizing radiation^89^, it seems highly likely that the increase of mitochondrial substitution rate by several hundreds of times compared to the mitochondrial substitution rate normally observed in amphibians have been caused by ionizing radiation. Nevertheless, this study does not allow specifying exactly the relationship between the artificial radionuclides exposure and the evolutionary processes estimated from genetic variations. The positive correlation between mitochondrial nucleotide diversity and ATDRs (currently ranging in the frogs samples at the CEZ from 0.007 to 22.4 μGy.h^-1^) is in agreement with an effect of ionizing radiation on genetic diversity, but there is no significant correlation between mitochondrial haplotype genetic diversity and ATDRs contrary to the results of Baker et al. on bank voles between haplotype genetic diversity and ambient dose rate^47^. The ATDR seems to be the most relevant dose rate estimator for a population over a time period, but it does not account for exposure of previous generations that occurred since the accident, even though possible transgenerational effects^90-92^ and evolutionary processes should be dependant of these historical doses. The measured mitochondrial substitutions may not only be caused by current exposure to artificial radionuclides, but may be also the result of mutations accumulated by individuals exposed to ionizing radiation in previous generations. There is no information on local tree frog population genetics before the accident and, thus, we cannot exclude uncertainties on the determination of the magnitude of the genetic modifications even if the use of Slavutych populations as a proxy of ancestral populations appears consistent. To fully understand the implication of ionizing radiation on the modification of the intensity of evolutionary processes, it should be valuable to compare these results with similar studies conducted in other radiocontaminated places like the Fukushima prefecture in Japan.

### The key role of mitochondrial DNA in evolutionary ecotoxicology

Our results show that the visible higher genetic diversity may not correspond to a classical evolutionary scenario (i.e. an ancestral population), and that mitochondrial markers are useful to assess the mutagenic effect of ionizing radiation^93^. Previous studies (e.g. Fuller et al.^94^) did not find any significant positive correlation between absorbed radiation (ATDRs ranging from 0.064 to 26.4 μGy.h^-1^) and nuclear genetic diversity in the freshwater crustacean *Asellus aquaticus* from the Chernobyl region. This study concludes that the exposure to ionizing radiation has not significantly influenced genetic diversity in *A. aquaticus* in the Chernobyl area. The analyses of mitochondrial markers might have provide other complementary information pointing towards a mutation process as showed in our study on *H. orientalis*. Measuring mitochondrial markers is thus important as a tool for estimate the modification of the intensity of evolutionary process, but also because of the probable consequences of mitochondrial mutations on individuals and populations. In humans, mtDNA mutations are responsible of several mitochondrial diseases like optic neuropathy^95^, MELAS^96^ and MERRF syndromes^97^. Because of the possible presence of different mtDNA in a single cell, disease symptoms associated with mtDNA mutation could be generated by quantitative changes in the proportion of mtDNA mutants^98^. Moreover, at the population level, the maternal transmission of mtDNA can prevent selection against mutations, which are deleterious only when expressed in males^99^ and can lead to a decrease in population viability^100^.

### The necessity of a large space and time scales

Genetic diversity can be sensible to many environmental parameters^50^ and considering a global phylogeographic context could help to overcome this issue. Examining only CEZ and Slavutych tree frogs populations would have been insufficient to draw reliable conclusions about evolutionary processes. However, by putting local estimations of genetic diversity of tree frogs (i.e. in the Chernobyl region) in a global phylogeographic context for the species, we were able to get a more accurate picture of the putative effects of radiocontamination on genetic variations and thus potential evolutionary processes of tree frogs populations in the CEZ. Our simulation data shows the need of a certain duration of exposure to radiation as well as the role of other factors like population size, generation time, and the mutation rate, to obtain a network pattern similar to that observed in the CEZ (Fig. 6d). It is possible that, depending on the life history of the organisms, genetic effects are different and/or not fully visible. Such difference might explain other recent findings showing an absence of visible radiation-induced mitochondrial microevolution^101^.

### Conclusions

Our study on the genetics of the Eastern tree frog populations in the CEZ suggests the existence of a strong mutation process on mitochondrial DNA, resulting in an unexpected genetic structure of the CEZ populations comparing to other European populations. One challenge now is to understand the possible consequences of this genotypic effect on population status. Due to the crucial role of mitochondria^102^ it seems unlikely these levels of mutation rate does not result in deleterious effects. The small population size predicted by our simulation may be a consequence of the elimination of non-viable individuals at birth, or due to other deleterious effects of ionizing radiation such as a reduction in breeding success (see e.g.^103^) or phenotypic disadvantage of mutations^104,105^. If the effects of these mutations do not fully compromise the maintenance of tree frog populations, it is not necessarily true for other organisms with different life history. With their large clutch sizes (up to 600 eggs per female per year^106^) tree frogs seem to be effective for supporting the deleterious effects of mutations, but it might not be the case for organisms with smaller litters for example. More detailed studies on species with different life history parameters are clearly needed to have a full picture of the eco-evolutionary effects of wildlife exposure to radioactive contamination.

## Materials and Methods

### 1 - Field work, capture and preparation of the samples

In May and June 2016, 2017, and 2018 during the breeding season, we collected a total of 216 *H. orientalis* individuals in 17 populations in wetlands located inside the CEZ and 2 outside the CEZ, i.e in the Slavutych region (Fig. 1b). For simplicity, we use here “population” in the meaning of “population sample”. These sites cover a gradient of ambient dose rates, that was measured using a hand-held radiometer (MKS-AT6130, ATOMTEX). The mean (±SD) ambient radiation dose rate varied from 0.044 to 32.4 μSv.h^-1^. After capture, individuals were kept in individual boxes with a perforated cover and 2 cm of water until the next morning when they were euthanized and dissected to sample tibia muscle. Collected tissue was quickly frozen at −196°C, transported to IRSN labs in Cadarache (France), and stored at −80 °C until DNA extraction. The geographic distances separating each pairwise combination of frog populations were estimated with ArcGIS and a UTM projection.

### 2 – Population-averaged dose rate calculation

The approach for population-averaged dose rate reconstruction was based on Giraudeau et al., 2018^63^ (See Supplementary note 1 for details). The two main differences compared with the protocol carried out by Giraudeau et al.^63^ are the radionuclides and the scenarios under consideration (Supplementary Fig. 3) because of the characteristics of the CEZ compared to the Fukushima situation. To summarize, soil activities (in Bq.kg^-1^) were extracted following Gashchak at al.^107^ from a spatial database using a geometric mean over a 400m radius area centred on each population location and using a time correction, and water activities were calculated using soil activities and distribution coefficients estimated for the Glubokoye lake^108^. In addition, frog activities (in Bq.kg^-1^) were estimated for each individual in femur bones for ^90^Sr, and in leg muscle for ^137^Cs in the IRL-SSRI Laboratory (Slavutych, Ukraine), and then reconstructed for the total frog knowing the total frog mass and the relative mass of bones (10%) and muscles (69%)^109^. A Canberra-Packard gamma-spectrometer with a high purity germanium (HPGe) detector (GC 3019) was used for measuring ^137^Cs sample activity concentrations and a Beta-spectrometer EXPRESS-01 was used for measuring ^90^Sr sample activity concentrations. For a more detailed description of measurement method of activity concentration see Beresford et al., 2020^110^. Then, dose coefficients (DCs) were calculated based on frog morphometry for internal exposure and four scenarios of external exposure using EDEN software^111^. DCs allow converting radionuclide activity (Bq.kg^-1^, Bq.L^-1^) into dose rate (μGy.h^-1^) and are specific for each radionuclide/scenario/organism combination. The total dose rate (in μGy.h^-1^) was calculated for each frog combining related dose coefficients and activities. Average total dose rates (ATDRs) were then obtained averaging for each population total dose rate of sampled individuals (Supplementary Fig. 4 and 5). Only the activity of ^90^Sr and ^137^Cs in frogs was measured, but the potential contribution of other less abundant radionuclides (^241^Am, ^238^Pu, ^239^Pu) to the total dose rate was estimated, leading to confirm their minor contribution to the total dose rate (on average less than a quarter of the total dose rate; see Supplementary Table 9, 10, 11 and Fig. 6). The total dose rate we assessed could potentially be slightly underestimated as other radionuclides than ^137^Cs and ^90^Sr were not included in the dose reconstruction. Nevertheless, in the CEZ the soil activity of ^90^Sr and ^137^Cs is correlated to the activity of other less abundant radionuclides^68^ as for the body activity of organisms such as small mammals^112^, thus our ATDR descriptor based on ^90^Sr and ^137^Cs is reliable for statistical tests.

### 3 - DNA extraction, sequencing and genotyping

DNA was extracted from tibia muscle using DNeasy Blood and Tissue Kit (Qiagen, Valencia, CA) following the manufacturer’s protocol. After the estimation of nucleotide concentration with a spectrophotometric measurement and an electrophoresis quality check, a 957 bp fragment of mitochondrial DNA, cytochrome b, and 21 nuclear microsatellites were studied (Supplementary Table 8). Mitochondrial and nuclear markers were used simultaneously in order to compare their different properties. To sequence the cytochrome b, a PCR amplification was performed using Hyla-L0 and Hyla-H1046 primers^62,113^. For each amplification session, a negative control was made using 3μL of water instead of extracted DNA, and an electrophoresis was done to control the proper functioning of the amplification. PCR-products were sequenced in both directions using Sanger sequencing (Eurofins, sequencing platform Cochin, France). The quality was checked using ab1 files. Sequences were aligned with MUSCLE program and corrected with MEGA^114^. In some cases, for the same position, an individual showed two different nucleotides. The mtDNA being haploid, it can be interpreted as an heteroplasmy^115^(i.e. the presence of multiple mtDNA haplotypes in an individual). For each of these individuals, the two haplotypes were considered. Four multiplex amplifications were then performed for the 21 microsatellite markers (Supplementary Table 8). Formamide and a Size Standard were added to the PCR-products and the whole was then genotyped with ABI-3100 Genetic Analyser (ThermoFisher Scientific). A second amplification and genotyping was carried out on 4 individuals in order to check the replicability of the method.

### 4 – Genetic analyses, mtDNA and nDNA

First of all, a quantitative analysis of population genetics was performed for the two types of markers. In order to avoid sample size artefacts, only populations with sample size higher or equal than 7 individuals were used to describe the genetic diversity. Because sample sizes were still non-homogeneous between populations, rarefaction technique was performed for cytochrome b to calculate haplotypic richness (nrH) and private estimated haplotype number (npH) using hp-rare^116^. The haplotype diversity (h)^117^, nucleotide diversity (π)^118^ and three estimators of θ index (θs, θk, θπ)^118-120^ were calculated for the cytochrome b using

ARLEQUIN^121^. To describe how the mitochondrial genetic variation is structured temporally and geographically, the Analysis of Molecular Variance (AMOVA)^69^ and the calculation of a differentiation index – pairwise Fst - were performed using ARLEQUIN. For microsatellites markers, we estimated the observed heterozygoty (Ho), the estimated heterozygoty under Hardy-Weinberg assumptions (He), the genetic diversity (Hs), the allelic richness (AR) and the private allelic richness (PA) using GENETIX^122^, ADZE^123^, and Fstat^124^. Pairwise Fst for microsatellites were calculated using Fstat. The absolute value of the lowest Fst for the mitochondrial and nuclear markers was added to every pairwise Fst in order to get only positive pairwise Fst. We calculated the ratio Fst/(1-Fst) in order to estimate genetic distance between populations, and represented these distances using Neighbour-Joining trees with genetic distances estimated with MEGA^114^.

A qualitative analysis of sequences was carried out for cytochrome b. The haplotypes were determined using DNAsp^125^, the haplotype network was calculated with the Median-Joining method^126^ and drawn using POPART^127^.

### 5 – Simulation of haplotype networks

Simulations of mitochondrial haplotype networks were conducted with a method close to the Approximate Bayesian Computation^128^ (for details on protocol see Supplementary note 2). Simulations of the haplotype evolution during 10 and 15 generations (assuming one generation respectively every two or three years^85,86^) in a unique population were conducted in R and Pegas library^129^ using different parameters: (i) the funder population size (N_0_) corresponding to specimens able to reproduce after the accident, (ii) the frequencies of haplotypes in the founder population based on the current diversity observed for the Slavutych populations, (iii) the population size for each year during the 10 or 15 generations (N_1-n_), (iv) the nucleotide substitution rate (*μ*) and (v) the number of generations. A haplotype network was generated for the last generation for each data set obtained with a set of value for prior parameters. After a first simulation running with classical wild frog population prior parameters^72^ (1000 simulations for each modality combination, a total of 6000 simulations), a second simulation was performed with different prior parameters calibrated from the first simulation results (100 simulations for each modality combination, a total of 21600 simulations) (see Supplementary Table 12 and 13 for prior parameters details). Each haplotype network was described by five statistics: the nucleotide diversity (π), the D Tajima, the haplotype richness (nrH), the haplotype diversity (h), and the number of steps separating the ancestral haplotype to the most distant haplotype plus one. A Principal Component Analysis (PCA) was performed to compare simulated and observed descriptive statistics. The two first principal component axes were used to visualise both sets of descriptive statistics. A Ward hierarchical cluster analysis on Euclidean distance was used to select the 5^th^ percentile of the simulated descriptive statistics closest to the observed descriptive statistics. The mean and median of the 5^th^ percentile simulated descriptive statistics were calculated to estimate the posterior parameters N_0_, the appropriate haplotype frequencies for the founder population, N_1-n_ and μ. Visualisation of the haplotype network for prior, 10_th_ and 15_th_ generation was done using TempNet^130^.

### 6 - Statistical analysis

Non-parametric Wilcoxon signed rank tests were performed to compare genetic diversity indices, between CEZ populations and outside CEZ populations, because of their non-normal distribution tested using Shapiro-Wilk test. Non-parametric Spearman rank test were used to test the correlation between genetic diversity indices and the population-averaged dose rate (ATDR). The correlation between two matrices, the genetic distance matrix obtained with the Fst linearization and the matrix of logarithm of geographical distance (in m), was performed using a Mantel test^70^ with the vegan package^131^. The part of pairwise population-averaged dose-rate differences between populations on the distances correlation was tested using a partial Mantel test. The significance of Mantel tests was estimated with 9999 permutations. All these tests were carried out on R version 3.6.1.^132^.

To test the demographical expansion hypothesis with haplotype networks, three statistical tests were performed on DNAsp. The neutral assumption of the absence of deviation from the mutation-drift equilibrium was tested using the Tajima’s D^133^ and Fu’s D*^134^. The distribution of pairwise differences between sequences was studied too, using the R2 statistic^135^.

## Acknowledgments

We are thankful to Jean-Michel Metivier (IRSN) for his help with GIS, to Yevgenii Gulyaichenko (Chernobyl Center, Slavutych, Ukraine) for his help in the field, to Andrii Maksymenko (Chernobyl Center, Slavutych, Ukraine) for the radionuclide assay of the samples, and to Julia Maklyuk and other staff of the Chernobyl Center (Slavutych, Ukraine) for the generic arrangement and logistic during the field work in Chernobyl. Field work in the Chernobyl Exclusion Zone was funded by EC EURATOM-60497 COMET project, Swedish Radiation Protection Agency-SSM (SSM2017-269 and SSM2018-2038) and Carl Tryggers Foundation (CT 16:344). Molecular analyses were funded by IRSN and the French NEEDS-Environnement grant. Radiation analyses were funded by Swedish Radiation Protection Agency-SSM (SSM2017-269 and SSM2018-2038). C. Car beneficed of an IRSN doctoral fellowship. P. Burraco was supported by a Carl Tryggers Foundation project CT 16:344 and by a Marie-Skłodowska-Curie individual fellowship (797879-METAGE project). G. Orizaola was supported by the Spanish Ministry of Science, Innovation and Universities “Ramón y Cajal” grant RYC-2016-20656.

