## Supplementary Information for "Unusual evolution of tree frog populations in the Chernobyl exclusion zone"

1 - Supplementary Figures
Supplementary Figure 1
Supplementary Figure 2
2 - Supplementary Tables
Supplementary Table 1
Supplementary Table 2
Supplementary Table 3
Supplementary Table 4
Supplementary Table 5
Supplementary Table 6
Supplementary Table 7
Supplementary Table 8
3 - Supplementary note 1: Total dose rate
3.1 – Scenarios
Supplementary Figure 3
3.2 – The distribution of total dose rate within and between populations
Supplementary Figure 4
Supplementary Figure 5
3.3 - Contribution of rare isotopes to the population-averaged dose rate
Supplementary Table 9
Supplementary Table 10
Supplementary Table 11
Supplementary Figure 6
4 – Supplementary note 2: Mitochondrial simulations
4.1 – Prior parameters
Supplementary Table 12
Supplementary Table 13
4.2 – Analysis of simulation results
4.2.1 - First simulation part
Supplementary Figure 7
4.2.2 - Second simulation part
Supplementary Figure 8
Supplementary Figure 9

1 – Supplementary Figures

Supplementary Figure 1: Example of heteroplasmy for the individual 12_9 of the A18 population for the 666 nucleotide position for Hyla-L0 and Hyla-H1046 primers.

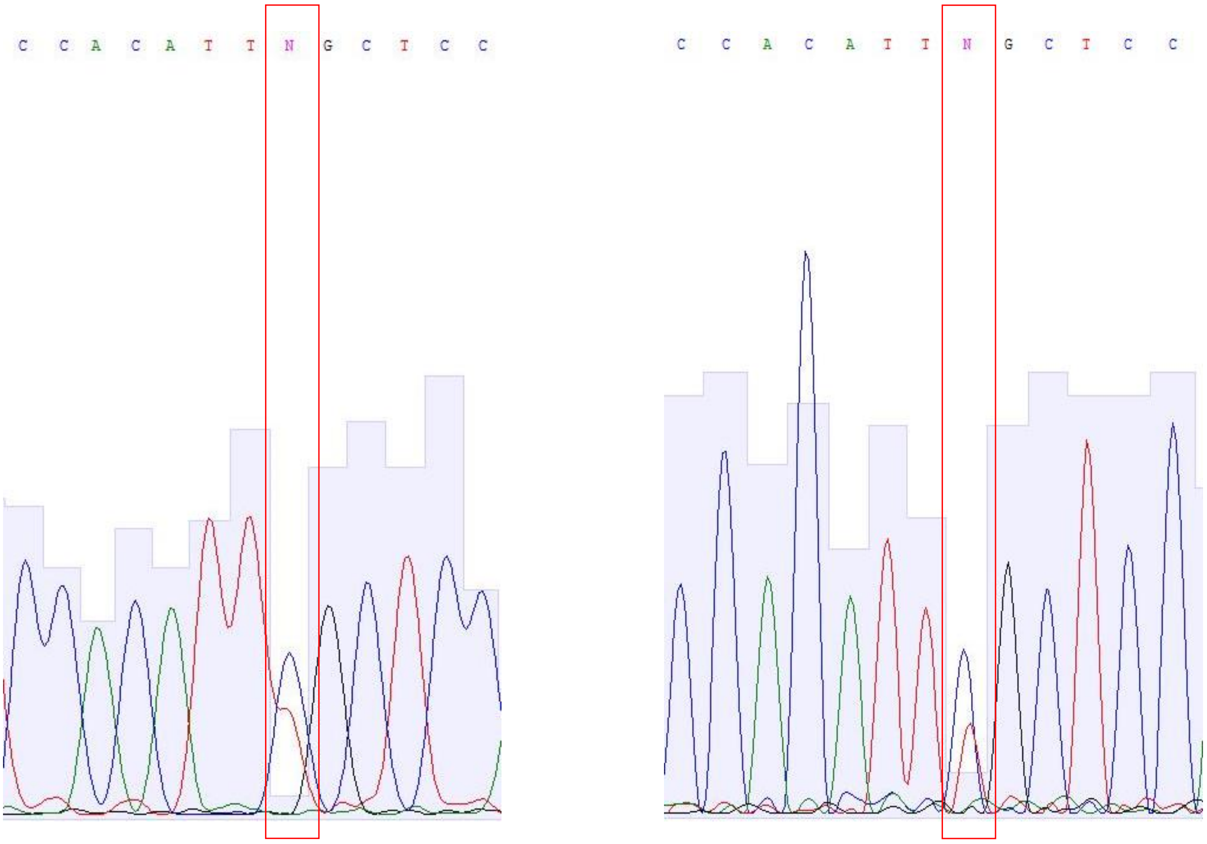

Supplementary Figure 2: Neighbor-Joining tree separating each year. a/b/c. From cytochrome b. d/e. From microsatellites. a/d. 2016. b/e. 2017. C. 2018

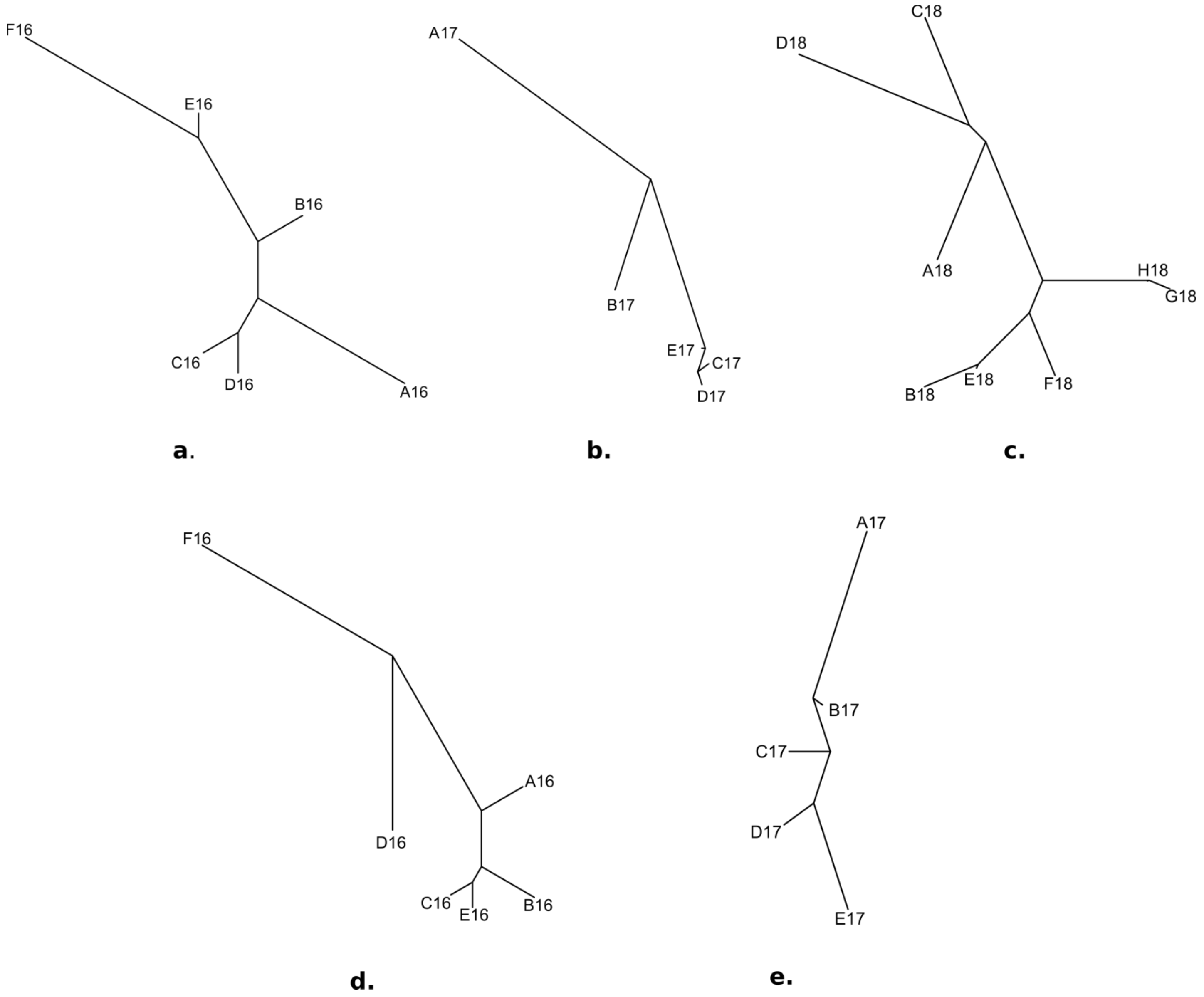

2 - Supplementary Tables

Supplementary Table 1: Haplotypes found at the Chernobyl exclusion zone (red), Slavutych (green) or both (orange). "." represents the same nucleotide as for the first haplotype for the position considered.

| Haplotypes | Nucleotide position | | | | | | | | | | | | | | | | | | | |
|---|---|---|---|---|---|---|---|---|---|---|---|---|---|---|---|---|---|---|---|---|
| | 57 | 78 | 92 | 96 | 105 | 151 | 153 | 199 | 259 | 279 | 322 | 426 | 427 | 594 | 666 | 690 | 806 | 874 | 934 | 939 |
| 1 | T | T | C | C | C | G | C | C | A | A | G | T | A | G | C | C | T | C | A | A |
| 2 | . | C | . | . | . | A | . | . | . | . | . | . | . | . | . | . | . | T | G | . |
| 3 | . | . | . | . | . | . | T | . | . | . | . | . | . | . | . | . | . | . | . | . |
| 4 | . | C | . | . | . | . | . | . | . | T | . | . | . | . | . | . | . | . | . | . |
| 5 | . | C | . | . | . | . | . | T | . | . | . | . | . | . | T | . | . | . | . | . |
| 6 | . | C | . | . | . | . | . | . | . | . | . | . | . | . | . | . | . | . | . | . |
| 7 | . | . | . | . | . | . | . | . | . | . | . | C | . | . | . | . | . | . | . | . |
| 8 | . | C | . | T | . | . | . | . | . | . | . | . | . | . | . | . | . | . | . | . |
| 9 | . | C | . | . | . | . | . | . | . | . | . | . | . | . | . | T | . | . | . | . |
| 10 | . | C | . | . | . | . | . | . | . | . | . | . | . | . | T | . | . | . | . | . |
| 11 | . | C | T | . | . | . | . | . | G | . | . | . | . | . | . | . | . | . | . | G |
| 12 | . | C | T | . | . | . | . | . | G | . | . | . | . | . | . | . | . | . | . | . |
| 13 | . | C | . | . | . | . | . | . | . | . | . | . | G | . | . | . | . | . | . | . |
| 14 | . | C | . | . | T | . | . | . | . | T | . | . | . | . | . | . | . | . | . | . |
| 15 | C | C | . | . | . | . | . | . | . | . | . | . | . | . | . | T | . | . | . | . |
| 16 | . | C | . | . | . | . | . | T | . | . | . | . | . | . | . | . | . | . | . | . |
| 17 | . | C | . | . | . | . | . | T | . | . | . | . | . | . | T | . | C | . | . | . |
| 18 | . | C | . | . | . | . | . | . | . | T | A | . | . | . | . | . | . | . | . | . |
| 19 | . | C | . | . | . | . | . | . | . | . | . | . | . | A | . | . | . | . | . | . |

Supplementary Table 2: Mutations measured in Chernobyl exclusion zone (red) and Slavutych (green) or both (orange) Oriental tree frogs on the $1^{st}$, $2^{nd}$ or $3^{rd}$ codon position, and modifications of amino acids.

| Position | 1st | 2nd | 3rd | AA |
|---|---|---|---|---|
| 57 | | | C/T | |
| 78 | | | C/T | |
| 92 | | C/T | | Ala/Val |
| 96 | | | C/T | |
| 105 | | | C/T | |
| 151 | A/G | | | Val/Ile |
| 153 | | | C/T | |
| 199 | C/T | | | |
| 259 | A/G | | | Met/Val |
| 279 | | | A/T | |
| 322 | A/G | | | Val/Ile |
| 426 | | | C/T | |
| 427 | A/G | | | Thr/Ala |
| 594 | | | G/A | |
| 666 | | | C/T | |
| 690 | | | C/T | |
| 806 | | T/C | | Ile/Thr |
| 874 | C/T | | | Leu/Phe |
| 934 | A/G | | | Thr/Ala |
| 939 | | | A/G | |

Supplementary Table 3: Populations and substitutions for the 7 tree frogs with heteroplasmy.

| Nucleotide position | Population | Number of individuals | Year of sampling |
|---|---|---|---|
| 153 – C/T | AB17 | 2 | 2017 |
| 199 – C/T | E17 | 1 | 2017 |
| 322 – C/T | F18 | 3 | 2018 |
| 666 – C/T | A18 | 1 | 2018 |

Supplementary Table 4: Statistical tests on different estimates of genetic diversity. Comparison of the genetic diversity of Chernobyl exclusion zone (CEZ) Oriental tree frog populations with other European populations (non-parametric Mann-Whitney/Wilcoxon tests for cytochrome b).

| Diversity index | W statistic | p.value | Median CEZ | Median other European populations |
|---|---|---|---|---|
| h | 91 | 0.005 ** | 0.7308 | 0.6071 |
| $\pi$ | 99 | 0.0004 *** | 0.0024 | 0.0008 |
| $\theta_S$ | 96 | 0.001 *** | 1.9868 | 0.8163 |
| $\theta_K$ | 92 | 0.006 ** | 2.5006 | 1.2535 |
| $\theta_\pi$ | 99 | 0.0004 *** | 2.183 | 0.7556 |
| nrH | 89 | 0.011 * | 3.140 | 2.520 |

Supplementary Table 5: Comparison of the genetic diversity of Chernobyl exclusion zone (CEZ) Oriental tree frog populations with other European populations (non-parametric Mann-Whitney/Wilcoxon tests for microsatellites markers).

| Diversity index | W statistic | p.value | Median CEZ | Median other European populations |
|---|---|---|---|---|
| He | 13 | 0.2398 | 0.2064 | 0.2400 |
| Ho | 17 | 0.5185 | 0.1887 | 0.2250 |
| Fis | 25 | 0.7972 | 0.0920 | 0.0670 |
| Gene diversity | 13 | 0.2398 | 0.2135 | 0.2410 |

Supplementary Table 6: Correlation tests between the genetic diversity of Oriental tree frog populations in the Chernobyl region, i.e. all the populations of Chernobyl exclusion zone and the two populations of Slavutych and the average total dose rates (ATDRs) of ionizing radiation absorbed by each population total (non-parametric Spearman rank correlation for cytochrome b).

| Diversity index | S statistic | rho | p.value |
|---|---|---|---|
| h | 658 | 0.1936 | 0.455 |
| $\pi$ | 294 | 0.6397 | 0.007 ** |
| $\theta_S$ | 536 | 0.343 | 0.1776 |
| $\theta_K$ | 581.86 | 0.287 | 0.264 |
| $\theta_\pi$ | 294 | 0.6397 | 0.007 ** |
| nrH | 634.78 | 0.222 | 0.3916 |

Supplementary Table 7: Non-parametric Spearman rank correlation for microsatellites markers between the genetic diversity of Oriental tree frog populations in the Chernobyl region, i.e. all the populations of Chernobyl exclusion zone and the two populations of Slavutych and the average total dose rates (ATDRs) of ionizing radiation absorbed by each population.

| Diversity index | S statistic | rho | p.value |
|---|---|---|---|
| He | 194 | -0.617 | 0.08573 |
| Ho | 148 | -0.233 | 0.6617 |
| Fis | 176 | -0.467 | 0.2125 |
| Gene diversity | 194 | -0.617 | 0.08573 |
| AR | 194.31 | -0.619 | 0.07535 |
| PA | 221.13 | -0.8427 | 0.004 ** |

Supplementary Table 8: Nuclear microsatellites used for genetic analysis.

| Primer (repeated pattern) | Forward primer | Reverse primer |
|---|---|---|
| Ha-T50 (CCG)7(CCA)1(CCT)5 | F: CAGCCCAACTGACTGGTTTT | R: GGGGAAGACTTTGACCCTCA |
| Ha-T53 (TT)1(TC)5 | F: TCTCCTGTCCTTCACCCAAC | R: CTTCCCAGCCTGGAACATC |
| Ha-T54 (AT)6 | F: GTGTGTAGGACCCAGGGAGA | R: TTGCTTCCGCTTGTGTAGTG |
| Ha-T55 (AG)7 | F: ATGGAAGGCTGAAGAGAGCA | R: CCAAAGGGTTAAATGCAGGA |
| Ha-T56 (AT)7 | F: TGCAAAAATGCCATGAAGTC | R: TTTGGAGACATCACGGTTGA |
| Ha-T58 (TCC)4…(TCA)6…(CTA)4 CTG(TCA)4 | F: TCCCGAAAGGACTACTGCTG | R: ACGCACAGGAGGAGAAAGAA |
| Ha-T60 (CAA)5(CAG)3 | F: ATTGCGAAAAACTGGTGGTT | R: GCTTTTCCCAGATCAACAGG |
| Ha-T61 (CAG)5 | F: CCGCAAAGATAATCCCAATC | R: AGGCTGCTGCATTAGATGGT |
| Ha-T63 (TTC)5 | F: TTCTGACCTCTCGGTTTGCT | R: ATGTAAAGGCGCTGATGGAG |
| Ha-T66 (CACACAT)1(TG)7 | F: CTCTTTCGGGTTCCATGCT | R: TCCATTGTGCTGATCGTGTT |
| Ha-T67 (CAT)6 | F: GGGCAGCTTTATTTTTCAGC | R: AGTGGCACCTCCAATAAAGG |
| Ha-T68 (CA)7 | F: AGGGCAGAGATACAGGCGTA | R: TGAAACAAATACCGGACTGC |
| Ha-A11 (CA)14 | F: CCTCCCTCACTCTGCTGAC | R: CAATCCCCGAAAAACATTG |
| Ha-A127 (TG)14 | F: CTCTGGGTTGCACTACTTAGTC | R: TTCAGGGCTAATTCTTTGTATG |
| Ha-B5R3 (TC)13 | F: CCCCTTTAGAGTCGCCATAC | R: AGCCATCTTGTGGTCAGTCA |
| WHA1-67 (CA)21 | F: GCTTTACACATGGGGGTAT | R: CACTCCTTTTAGAGTATGTTGTTG |
| Ha-D104 (TAGA)7 | F: GCTGGCTGACTTATTCTTTG | R: TCTTCTCTCCACGGTCTTC |
| Ha-D115 (TAGA)16 | F: GTTTTTCGATTCCTGGATAAC | R: TGGGAGTTTTCAAAAGTGAC |
| Ha-E2 (CAA)7 | F: ACAACTTCCAACTGGAGTCAAC | R: CCTTAGTGGGAGCTGTAATCAC |
| Ha-A110 (CA)15 | F: AAGGGTTAAATCACCTATCC | R: ACGCAAAAAACATCTGTG |
| Ha-A119 (CT)14(CA)6TA(CA)14 | F: CAACTTCCCCCTCTGTTC | R: GCTGAGTGTGAGTGTGTTTG |

3 - Supplementary note 1: Total dose rate

3.1 – Scenarios

Supplementary Figure 3: Exposure scenarios applied to estimate the Dose Coefficients (illustration modified from Giraudeau et al, 2018[1]). Because of the characteristics of the Chernobyl exclusion zone compared to the Fukushima situation, the depth of the different microhabitats and the time of exposure to the four scenarios has been modified.

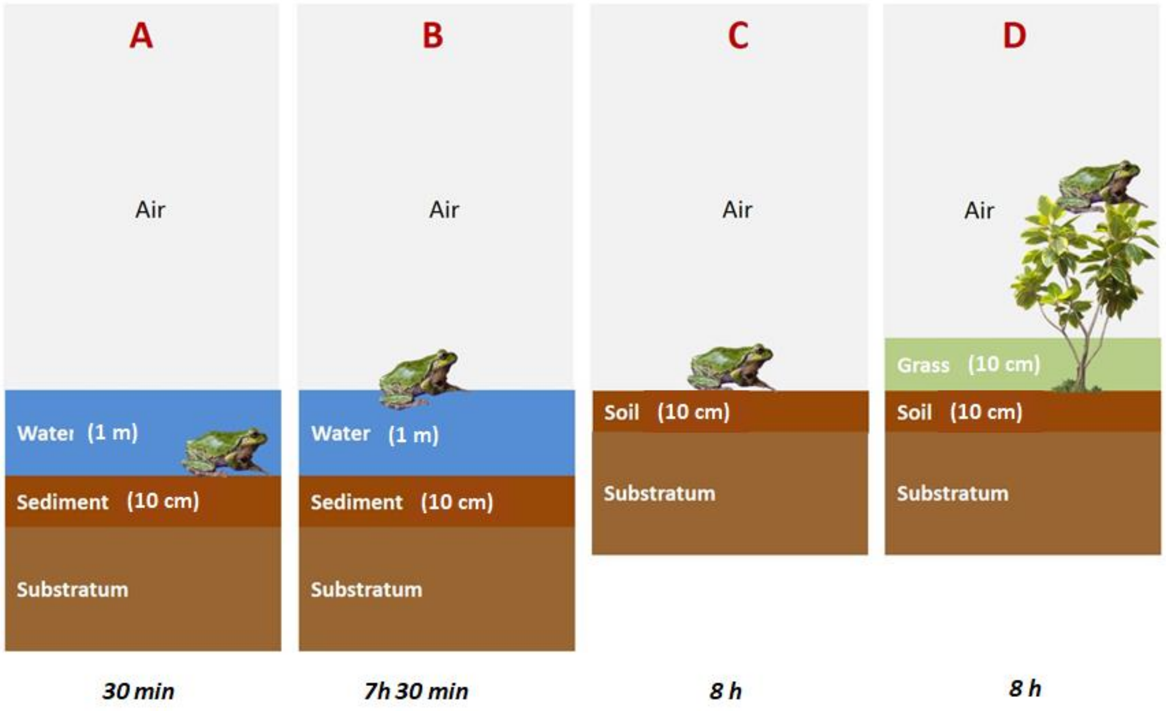

3.2 – The distribution of total dose rate within and between populations

Supplementary Figure 4: Dose distribution within and between populations. Representation of total dose rates absorbed by individual (TDRs) (µGy.h$^{-1}$) for the Oriental tree frog populations of the Chernobyl exclusion zone. Colours represent the ambient dose rate gradient measured on the field (>10 µGy.h$^{-1}$, dark red; >5 µGy.h$^{-1}$, red, >3 µGy.h$^{-1}$, orange, >2 µGy.h$^{-1}$, dark yellow; >1 µGy.h$^{-1}$, yellow; <1 µGy.h$^{-1}$, blue).

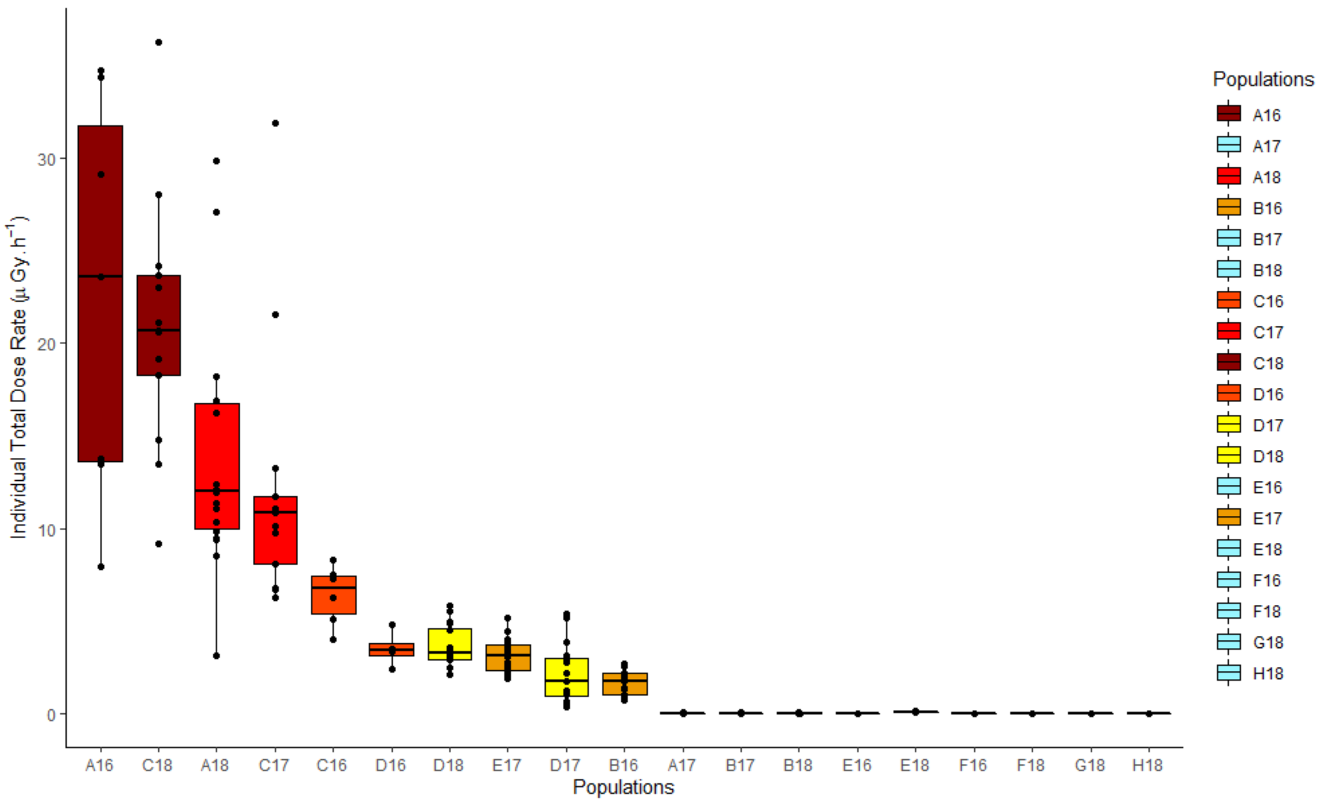

Supplementary Figure 5: Representation of the internal (blue) and external (pink) average total dose rates (ATDRs) of ionizing radiation absorbed (μGy.h$^{-1}$) for Chernobyl exclusion zone populations rate.

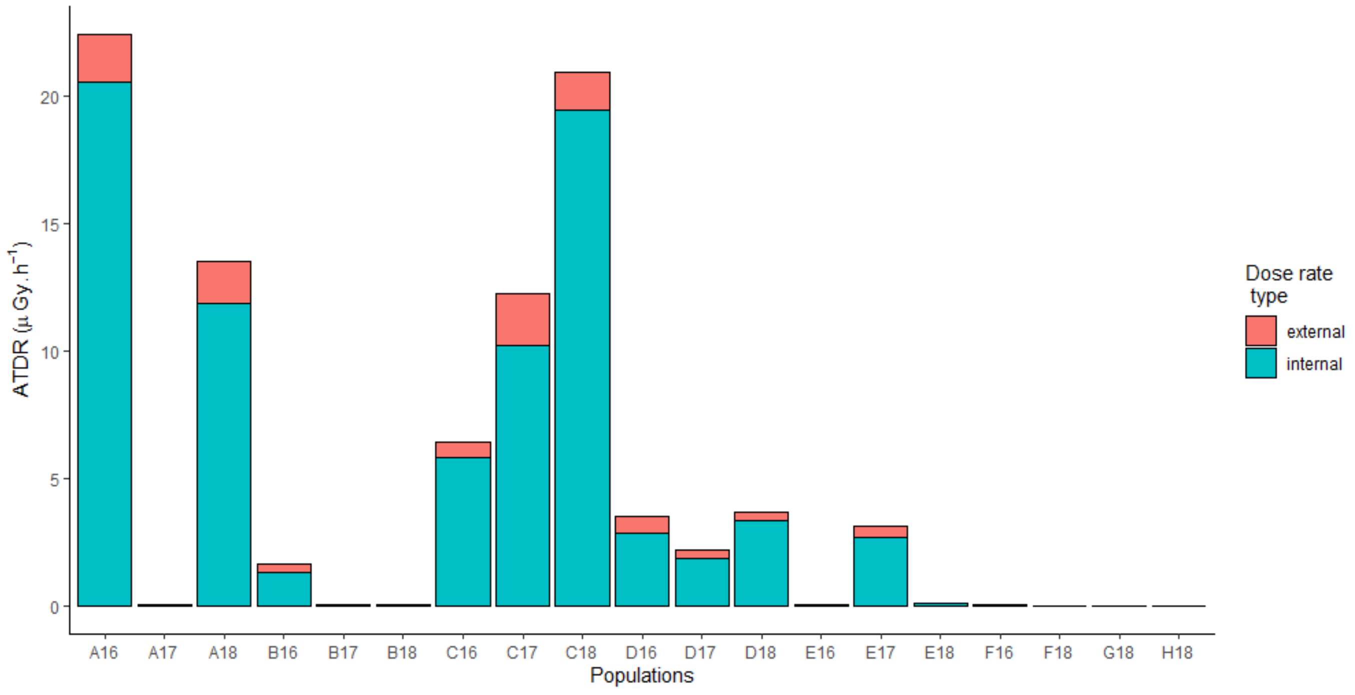

3.3 - Contribution of rare isotopes to the population-averaged dose rate

In order to assess the contribution of rare radionuclides compared to $^{137}Cs$ and $^{90}Sr$ on the total dose rate, dose rate relative to each radionuclide was reconstructed. A set of realistic extreme activity concentrations in soils estimated for February 2009 for every radionuclide was used[2–4]. The radioactive decay was applied to each minimum and maximum value.

Supplementary Table 9: Minimal and maximal soil activity concentration ($Bq.kg^{-1}$) in 2017 for radionuclides present in the Chernobyl exclusion zone.

| | Soil activity concentration ($Bq.kg^{-1}$) | |
|---|---|---|
| | Minimum | Maximum |
| $^{137}Cs$ | 56.3 | 395942 |
| $^{90}Sr$ | 29.7 | 368695 |
| $^{241}Am$ | 0.67 | 12192 |
| $^{238}Pu$ | 0.76 | 3435 |
| $^{239}Pu$ | 2.43 | 8761 |
| $^{240}Pu$ | 2.43 | 8753 |
| $^{234}U$ | 0.95 | 20.0 |
| $^{238}U$ | 1.07 | 11.0 |
| $^{60}Co$ | 0.04 | 196 |
| $^{154}Eu$ | 0.51 | 1130 |

The internal activity concentration was reconstructed using concentration ratio estimated by Beresford et al.[5] on *Rana arvalis* and ERICA 1.3 [6] for other non-documented radionuclides.

Supplementary Table 10: Minimal and maximal frog activity concentration ($Bq.kg^{-1}$) in 2017 for radionuclides present in the Chernobyl exclusion zone.

| | Frog activity concentration ($Bq.kg^{-1}$) | |
|---|---|---|
| | Minimum | Maximum |
| $^{137}Cs$ | 21.88 | 154021 |
| $^{90}Sr$ | 19.03 | 235964 |
| $^{241}Am$ | 0.00 | 10.75 |
| $^{238}Pu$ | 0.02 | 88.28 |
| $^{239}Pu$ | 0.06 | 225.16 |
| $^{240}Pu$ | 0.06 | 224.96 |
| $^{234}U$ | 0.00 | 0.10 |
| $^{238}U$ | 0.01 | 0.05 |
| $^{60}Co$ | 0.01 | 37.39 |
| $^{154}Eu$ | 0.02 | 38.44 |

Dose coefficient for each radionuclide/scenario was calculated with EDEN software [7]. Similarly to the total dose rate calculation presented on Material and Methods, external and internal dose rate were compiled to estimate the total dose rate for minimal and maximal soil activity concentration.

Supplementary Table 11: Minimal and maximal Total dose rate ($\mu Gy.h^{-1}$) for radionuclides present in the Chernobyl exclusion zone.

| | Total dose rate | |
|---|---|---|
| | Minimum | Maximum |
| $^{137}Cs$ | 0.003 | 20.4 |
| $^{90}Sr$ | 0.007 | 80.3 |
| $^{241}Am$ | 0.00002 | 0.42 |
| $^{238}Pu$ | 0.0006 | 2.56 |
| $^{239}Pu$ | 0.002 | 6.12 |
| $^{240}Pu$ | 0.002 | 6.13 |
| $^{234}U$ | 0.0001 | 0.003 |
| $^{238}U$ | 0.0001 | 0.001 |
| $^{60}Co$ | 0.000004 | 0.02 |
| $^{154}Eu$ | 0.00002 | 0.05 |

Supplementary Figure 6: Distribution of radionuclide dose rate among total dose rate for minimal and maximal soil activity concentration. 90Sr and 137Cs are the principal contributors of total dose rate (respectively 69% and 87%).

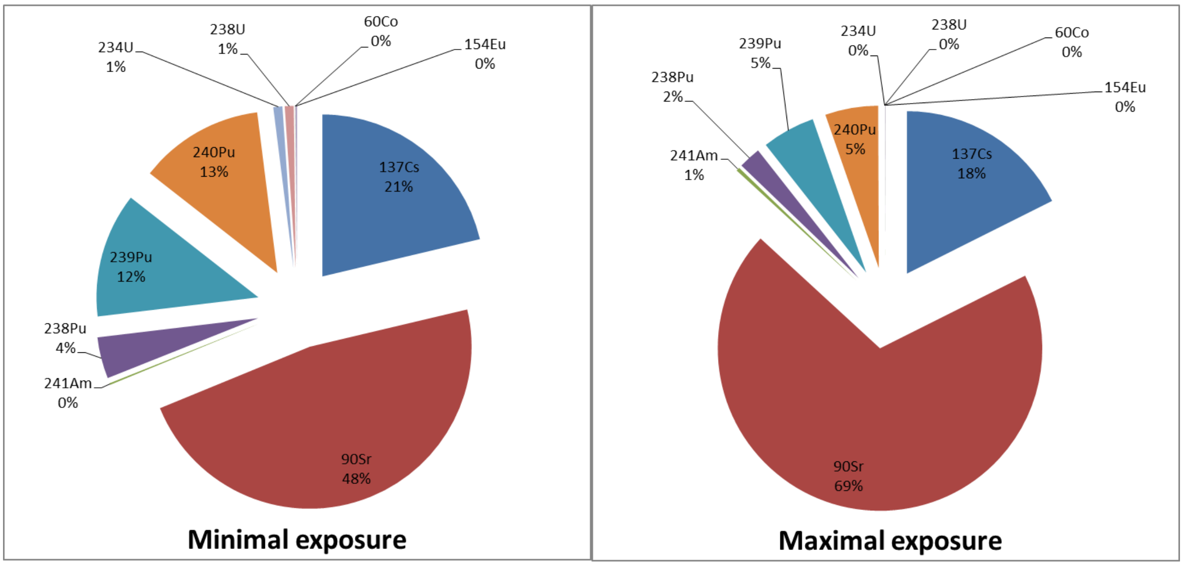


1. Giraudeau, M. *et al.* Carotenoid distribution in wild Japanese tree frogs (*Hyla japonica*) exposed to ionizing radiation in Fukushima. *Sci. Rep.* **8**, 1–11 (2018).
2. Theodorakopoulos, N. Analyse de la biodiversité bactérienne d’un sol contaminé de la zone d’exclusion de Tchernobyl et caractérisation de l’interaction engagée par une souche de Microbacterium avec l’uranium. (Université d’Aix-Marseille, 2013).
3. Chapon, V. *et al.* Microbial diversity in contaminated soils along the T22 trench of the Chernobyl experimental platform. *Appl. Geochem.* **27**, 1375–1383 (2012).
4. Lecomte-Pradines, C. *et al.* Soil nematode assemblages as bioindicators of radiation impact in the Chernobyl Exclusion Zone. *Sci. Total Environ.* **490**, 161–170 (2014).
5. Beresford, N. A. *et al.* Radionuclide transfer to wildlife at a ‘Reference site’ in the Chernobyl Exclusion Zone and resultant radiation exposures. *J. Environ. Radioact.* **211**, 105661 (2020).

6. Brown, J. E. *et al.* The ERICA Tool. *J. Environ. Radioact.* **99**, 1371–1383 (2008).
7. Beaugelin-Seiller, K., Jasserand, F., Garnier-Laplace, J. & Gariel, J. C. Modeling radiological dose in non-human species: principles, computerization, and application. *Health Phys.* **90**, 485–493 (2006).

### 4 - Supplementary note 2: Mitochondrial simulations

#### 4.1 – Prior parameters

Supplementary Table 12: Prior parameters of the first simulation part. These parameters are chosen to be representative of a classical frog population in a wild environment.

| | |
|---|---|
| Founder population size ($N_0$) | $N_0 = 500$ |
| Frequencies of haplotypes in the founder population | Two populations were sampled in Slavutych and may represent the closest local control population. The two set of haplotype frequencies were chosen as two modalities for this parameter. G18 (4 haplotypes): f1 = 0.6, f2 = 0.2, f3 = 0.1, f4 = 0.1 and H18 (2 haplotypes): f1 = 0.95, f2 = 0.05, f3 = 0.0, f4 = 0.0. Each haplotype was separated by one mutation. |
| Population size for each generation ($N_{1-n}$) | For each generation, $N_{min}$ and $N_{max}$ were sampled in a uniform distribution U(1000-5000) corresponding to an increase of the population size for the first generation $N_1$ and then a fluctuating population size in the CEZ (balance between the number of dead and alive specimens). |
| Generation time | Because of the 30 years separating us from the Chernobyl nuclear power plant accident, a number 15 and 10 tree frog generations was chosen, considering a generation time of respectively 2 and 3 years[1,2]. |
| Nucleotide substitution rate $\mu$ | A classical rate of nucleotide substitution in mitochondrial DNA for amphibian equal to $20.37 \times 10^{-9}$ substitution/nucleotide/year (in Lynch 2007 p. 320[1]) was chosen. Considering that one generation for amphibian equal to one year according to Lynch 2007 and considering the 950 bases of the cytochrome b gene we obtain a nucleotide substitution rate μ of $1.94 \times 10^{-4}$ substitution/haplotype/year (≈ 0.0002). |

Supplementary Table 13: Prior parameters of the second simulation part. These parameters are chosen considering the results of the first simulation part: the first parameters being not able to obtain the diversity of populations of Chernobyl exclusion zone, population sizes should be smaller and nucleotide substitution rate should be higher.

| | |
|---|---|
| Founder population size ($N_0$) | Three modalities: $N_0$ = 100, 250 or 500. |
| Frequencies of haplotypes in the founder population | As for the first simulation part, the two set of haplotype frequencies of Slavutych populations were chosen as two modalities for this parameter. G18 (4 haplotypes): f1 = 0.6, f2 = 0.2, f3 = 0.1, f4 = 0.1 and H18 (2 haplotypes): f1 = 0.95, f2 = 0.05, f3 = 0.0, f4 = 0.0. Each haplotype was separated by one mutation. |
| Population size for each year ($N_{1\text{-}30}$) | Three modalities: for each year, $N_{min}$ and $N_{max}$ were sampled in a uniform distribution U(50-100), U(100-200) or U(200-300). |
| Generation time | Because of the 30 years separating us from the Chernobyl nuclear power plant accident, a number of 15 and 10 tree frog generations was chosen, considering a generation time of respectively 2 and 3 years[1,2] |
| Nucleotide substitution rate $\mu$ | Six modalities: 0.005, 0.01, 0.02, 0.04, 0.06, 0.08 using an infinite site model. |


1. Altunisik, A. & Özdemir, N. Body size and age structure of a highland population of *Hyla orientalis* Bedriaga, 1890, in northern Turkey. *Herpetozoa* **26**, 49–55 (2013).
2. Özdemir, N. *et al.* Variation in body size and age structure among three Turkish populations of the treefrog *Hyla arborea*. *Amphibia-Reptilia* **33**, 25–35 (2012).
3. Lynch, M. & Walsh, B. *The origins of genome architecture*. vol. 98 (2007).

4.2 – Analysis of simulation results

Supplementary Figure 7: Distribution of the five haplotype network descriptive statistics for the five percentile closest simulated values. The simulated median (red) and mean (blue) present huge differences from the observed (green, Supplementary Table 4).

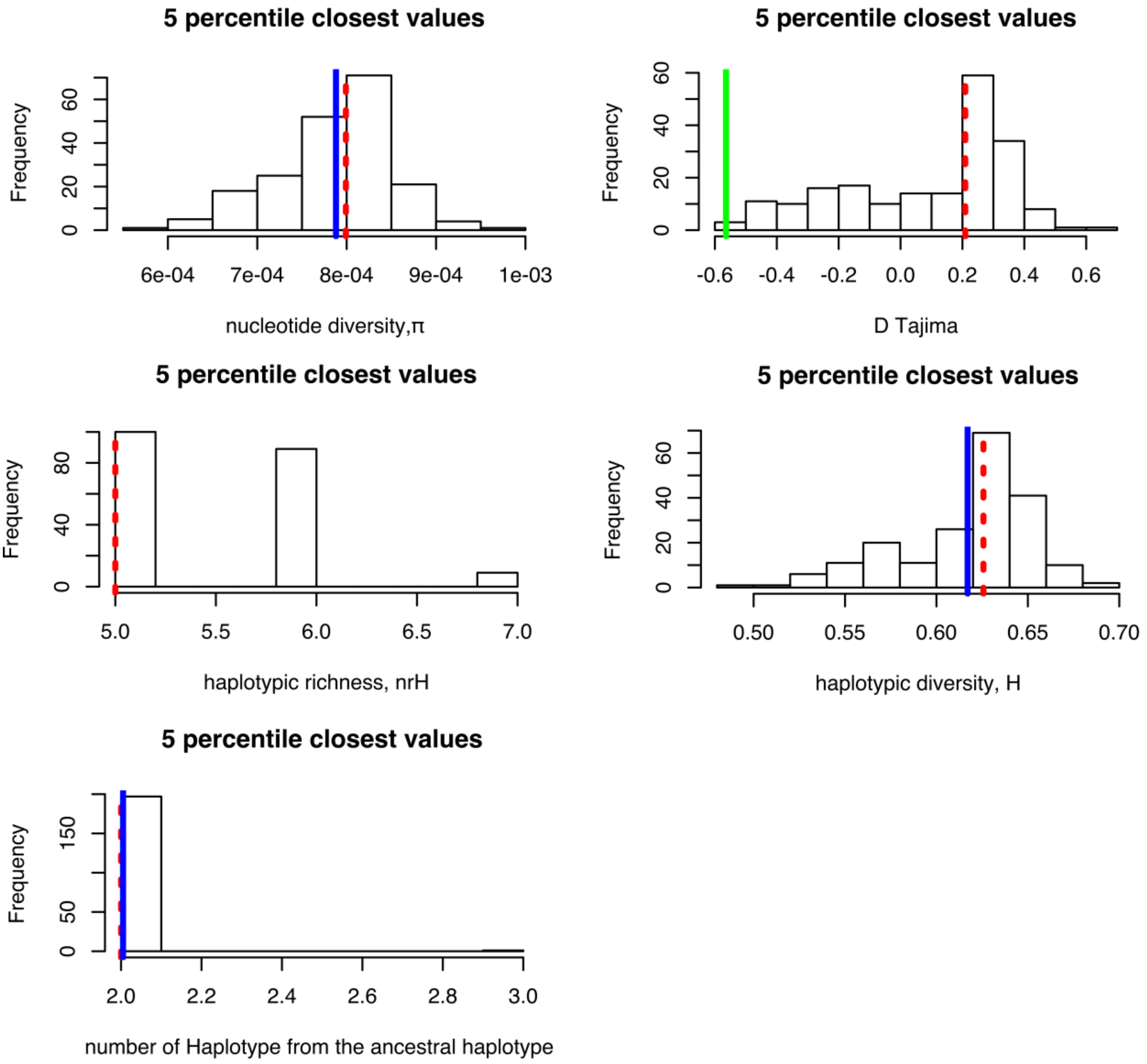

4.2.2 - Second simulation part

Supplementary Figure 8: Distribution of the values of parameters $\mu$, $N_{max}$, $N_0$, number of generations and haplotype frequencies of the founder population for the 5 percentile closest haplotype networks (red = median, blue = mean).

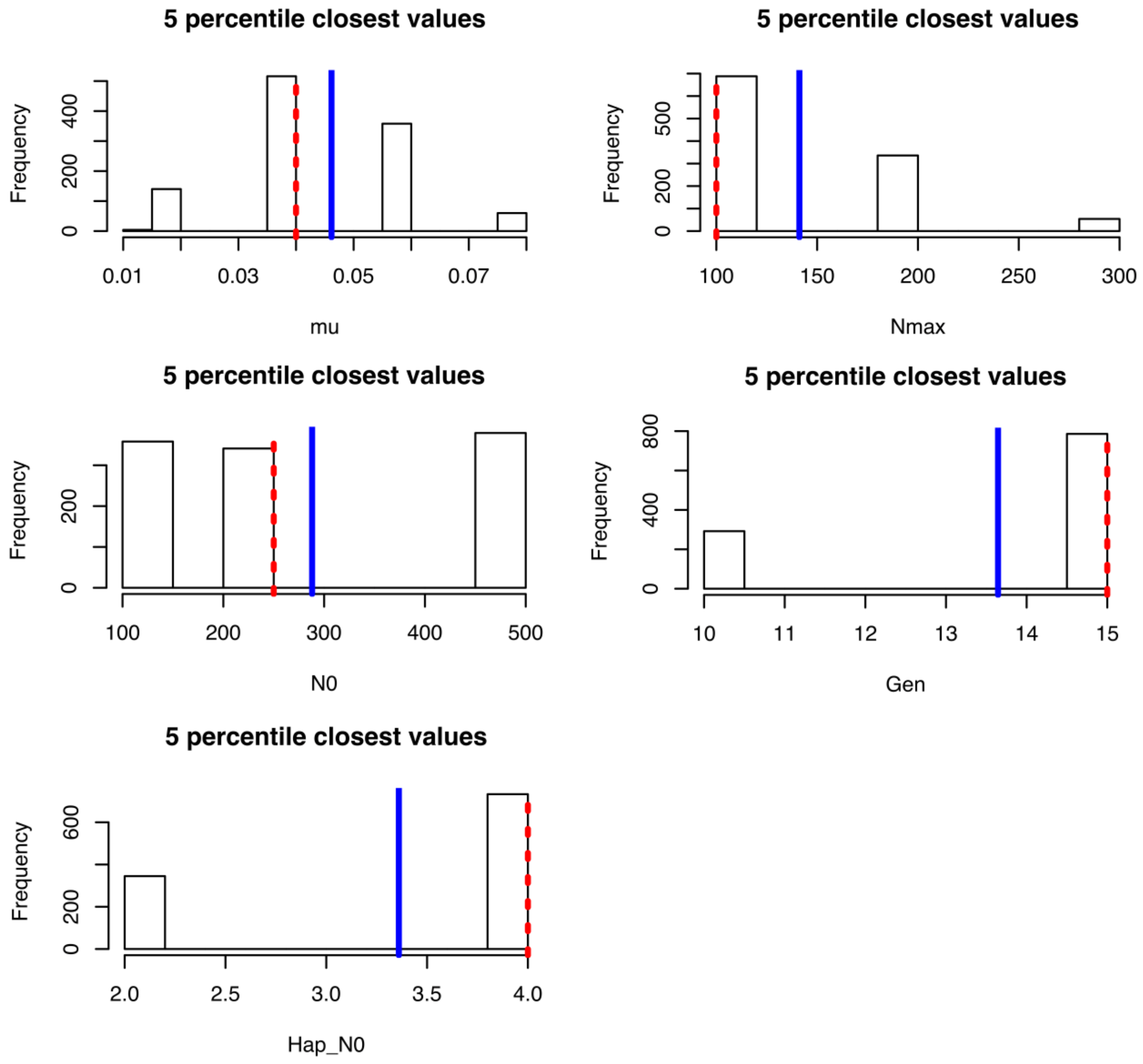

Supplementary Figure 9: Distribution of the five haplotype network descriptive statistics for the five percentile closest simulated values. The simulated median (red) and mean (blue) are close to the observed ones (green = mean, Supplementary Table 4).

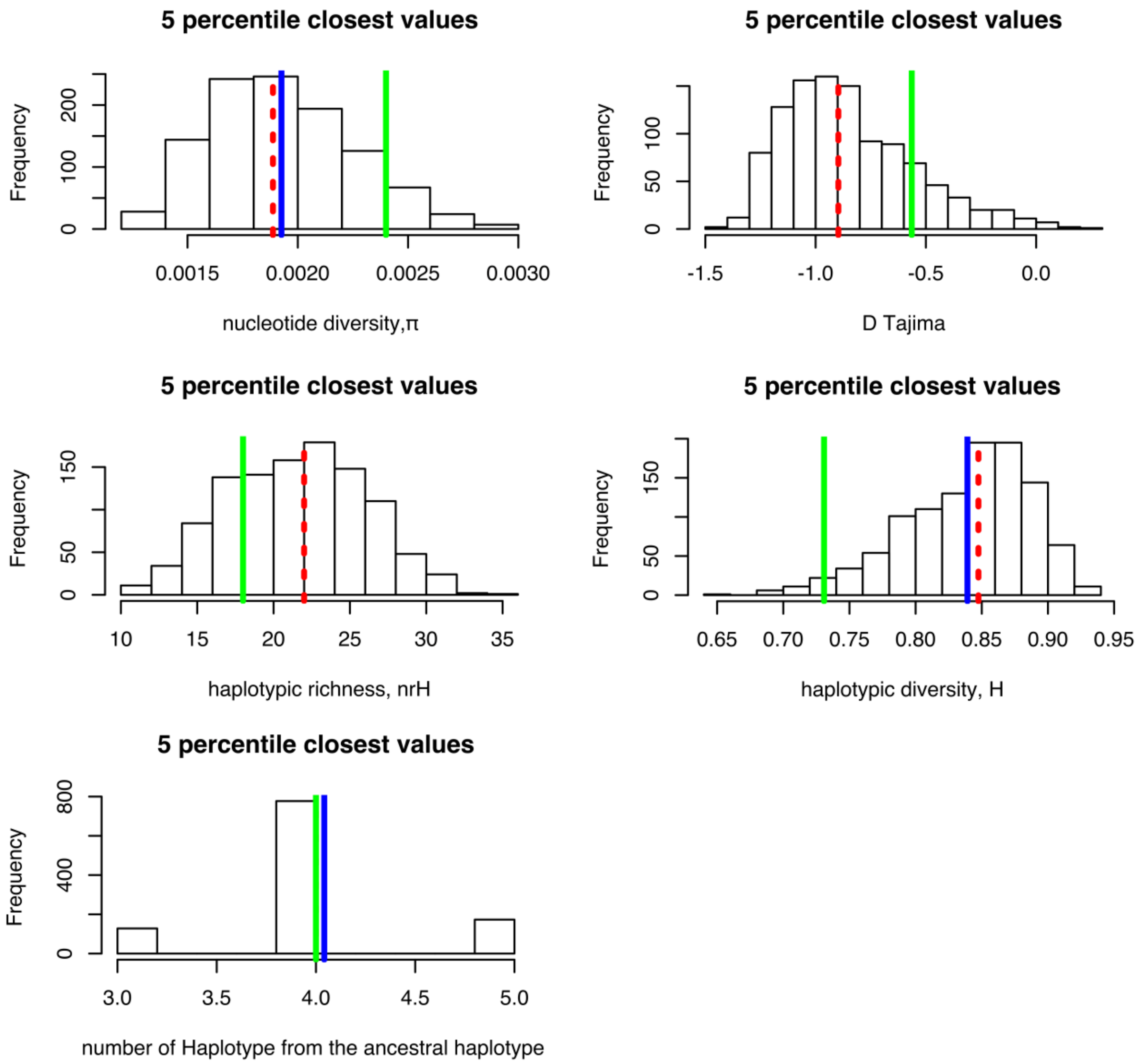
